# Reproducible growth of *Brachypodium distachyon* in fabricated ecosystems (EcoFAB 2.0) reveals that nitrogen form and starvation modulate root exudation

**DOI:** 10.1101/2023.01.18.524647

**Authors:** Vlastimil Novak, Peter F. Andeer, Benjamin P. Bowen, Yezhang Ding, Kateryna Zhalnina, Connor Tomaka, Amber N. Golini, Suzanne M. Kosina, Trent R. Northen

## Abstract

Understanding plant-microbe interactions requires examination of root exudation under nutrient stress using standardized and reproducible experimental systems. We grew *Brachypodium distachyon* hydroponically in novel fabricated ecosystem devices (EcoFAB 2.0) under three inorganic nitrogen forms (NO_3_^−^, NH_4_^+^, NH_4_NO_3_), followed by nitrogen starvation. Analyses of exudates with LC-MS/MS, biomass, medium pH, and nitrogen uptake showed EcoFAB 2.0’s low intra-treatment data variability. Furthermore, the three inorganic nitrogen forms caused differential exudation, generalized by abundant amino acids/peptides and alkaloids. Comparatively, N-deficiency decreased N-containing compounds but increased shikimates/phenylpropanoids. Subsequent bioassays with two shikimates/phenylpropanoids (shikimic and *p*-coumaric acids) on the rhizobacterium *Pseudomonas putida* or *Brachypodium* seedlings revealed that shikimic acid promoted bacterial and root growth, while *p*-coumaric acid stunted seedlings. Our results suggest: (i) *Brachypodium* alters exudation in response to nitrogen status, which can affect rhizobacterial growth; and (ii) EcoFAB 2.0 is a valuable standardized plant research tool.

**Teaser:** EcoFAB 2.0, a novel fabricated ecosystem device, has low data variability in studies of plant traits.

## Introduction

Plants exude up to 40% of assimilated carbon from roots into the soil, primarily as diverse small compounds such as sugars, amino acids, organic acids, and secondary metabolites (*1*). These chemical cocktails are dynamic, complex, and associated with diverse functions in the rhizosphere (*2, 3*). Some exudate metabolites, particularly cinnamic acid derivatives (e.g., *p*-coumaric, caffeic, or ferulic acids), were shown to have allelopathic effects in plants (*4, 5*). In contrast, some exogenous metabolites can promote root growth, significantly contributing to plant C budget (*6*). Exudates can also inhibit the activity of nitrifying bacteria, while other compounds support N_2_-fixing symbiotic microorganisms (*7*). Since these compounds generate a nutrient-rich region in soils, they are a primary driver for microbial community formation that ideally benefit the plant (*8*). For example, specific root exudates from the wild oat *Avena barbata*, including several aromatic acids (nicotinic, shikimic, salicylic, cinnamic, and indole-3-acetic) were shown to be preferentially consumed by rhizosphere bacteria (*3*). Plant exudation can also alter the cycling of soil organic matter to bioavailable N in soil (*9*). In particular, oxalic acid can cause the microbe-independent destabilization of mineral-associated organic N, while glucose can activate microbial-mediated N-mining (*10, 11*). In perspective, the optimized N cycling via plant-microbe interactions can be a key to the increased production of bioenergy grasses on marginal N-limited soils (*12, 13*).

Nitrogen is an essential and often plant growth-limiting macronutrient that plants uptake through roots primarily as ammonium (NH_4_^+^) and nitrate (NO_3_^−^) and to a lesser extent as organic-N (*6, 14, 15*). Previous studies showed that inorganic nitrogen (iN) form affects plant development. For example, the supply of NH_4_^+^ or NO_3_^−^ alters the phenotypes and cell wall composition of the model grass plant *Brachypodium distachyon* (*16*). On the other hand, N deficiency causes changes in the transcriptome, chlorosis due to chlorophyll degradation, a decrease in plant foliar N content, and changes in auxin levels resulting in initial root foraging (*14, 17–19*). Multiple studies showed exudate changes by N supply in several plant grass species. For example, a recent paper showed the increase of aromatic and other organic acids in marginal N-depleted soils of the switchgrass rhizosphere, while added nitrogen (+N) and nitrogen phosphate (+NP) fertilizer increased the abundance of amino acids (*20*). A study on rice (*Oryza sativa*) found that roots actively release many metabolites in response to N and P deficiency (*21*). Another study showed that axenically grown maize in N, P, Fe, or K deficient conditions changed the abundance of specific exuded amino acids, sugars, and organic acids (*22*).

The annual grass *Brachypodium distachyon*, an accepted model for many bioenergy and grain crops (23), has many favorable traits for laboratory studies, including a relatively small diploid genome, small size, and a short reproduction cycle (*23*). Despite the importance of *B. distachyon* to understanding fundamental processes associated with grass growth and genetics, currently, there is a lack of studies systematically exploring the effects of inorganic N (NH_4_NO_3_, NO_3_^−^, or NH_4_^+^) starvation on root exudation in *B. distachyon*. Given that changes in response to the form of inorganic nitrogen (iN) may be subtle, we require methods for achieving highly reproducible and sterile plant growth, exudate collection, and phenotyping across hundreds of plants. We have previously described ‘EcoFAB’ devices that were shown to support reproducible *B. distachyon* growth and phenotyping across 4 different laboratories (*24*). However, these EcoFABs were made by ‘hand’ molding polydimethylsiloxane rubber in 3D printed molds followed by chemically bonding them to microscope slides, thus limiting the scale of studies they can support (*25*).

Here, we aim to increase the understanding of *B. distachyon* exudation dynamics under varying N supply. We hypothesize that iN form and starvation will change the plant exudate profile and that iN-starvation-induced exudates have diverse functions in the rhizosphere that will alter the growth of microbes and seedlings. To test this hypothesis, we developed and used a new EcoFAB device (EcoFAB 2.0), made using a high-throughput fabrication. The combination of different inorganic N sufficient (iN+) treatments (NH_4_NO_3_, NO_3_^−^, or NH_4_^+^) with N starvation (iN-) in a hydroponic environment of EcoFAB 2.0 devices allowed us to assess the modulation of root exudation in *B. distachyon* Bd21-3 via LC-MS/MS-based metabolomics analysis followed by spectral matching using an online database as well as feature-based molecular networking. We combined the LC-MS/MS analysis with the Global Natural Products Social Molecular Networking (GNPS) tool, which provides us with feature annotations based on similarities to spectral libraries (*26*). We also used NPClassifier, a deep-neural network tool that classifies compounds based on structural characteristics (*27*). Metabolomics was followed by plant and bacterial bioassays to access the functions of selected exudate compounds.

## Results

### EcoFAB 2.0 development

The EcoFAB (https://eco-fab.org/) platform aims to create standardized systems to study environmental microbiomes and plant exudation (*28*). Initial devices were made using polydimethylsiloxane rubber and 3D-printed molds (*25*). At the same time, we found that the resulting devices had high cross-laboratory reproducibility of plant morphology and exometabolome (*24*). EcoFAB 2.0 is another sterile, self-contained system suitable for sterile plant growth and imaging (U.S. Patent Application No: 16/876,415). Version 2.0 is larger to allow for studying small plants like *B. distachyon* over their entire life cycle or large plants but only at early growth stages. Critically, EcoFAB 2.0 is made using injection molding, which enables high throughput fabrication. The EcoFAB 2.0 comprises 3 injection molded polycarbonate parts: a silicone gasket, a glass slide, and 8 screws (**Protocol S1**). It has a circa 10 mL root chamber that slightly slopes downward away from the plant port while the device is sitting flat and has two sampling ports and a 264 mL shoot chamber (height x length x width = 50 × 66 × 80 mm) with 4 luer-compatible gas vent ports (**Fig. S1A**). The root zone used in this study was c.a. 3 mm thick. However, this thickness is determined by the gasket thickness and is, therefore, adjustable. All parts can be sterilized by autoclaving or ethylene oxide (*29*). The base dimensions of the EcoFAB 2.0 are the same as a standard multiwell plate (SBS format), making them compatible with liquid handling equipment (**Fig. S1A**).

EcoFABs 2.0 are compatible with plant hydroponic and soil experiments (**Fig. S1B**) and have been used to grow not only *B. distachyon* but also *Arabidopsis thaliana, Lotus japonicus*, and *Camelina Suneson* for at least 4 weeks. Because the parts are made of injection-molded polycarbonate, dyes can be added to the EcoFAB base to shield the roots chamber from specific wavelengths or all visible light for light-sensitive rhizosphere experiments (**Fig. S1B**). During development, we have also tested visualization of root architecture by scanning with flatbed scanners, inverted microscopy on roots, sampling of microbes for determination of root microbiome structure by 16S sequencing, metabolomics on spent medium, and gas analysis in the shoot top chamber (**Fig. S1C**). While large glass slides of standard thickness were used for this study, the backing plate in the EcoFABs can be reversed to accommodate thinner slides (i.e., 0.55 mm Gorilla Glass). In our hydroponic experiments, we tested the sterility of EcoFAB 2.0 at the end of weeks 3 and 5 by incubating the spent plant growth media from each replicate on Luria-Bertani (LB) agar plates for several days at 27 °C in the dark; this revealed 100% sterility (not shown) of the EcoFAB 2.0 system over the experimental period of 5 weeks.

### Form of inorganic nitrogen changes plant biomass

We grew *B. distachyon* in a hydroponic EcoFAB 2.0 setup and analyzed plant biomass after 5 weeks (**Fig. 1A and B**). The type of iN treatment was correlated with phenotypical differences in the *B. distachyon* Bd21-3 plants (**Fig. 1C)**. After 5 weeks, all plants were at the tillering/main stem elongation stage (**Fig. 1C**). The two weeks on iN-resulted in mild chlorosis on the bottom leaves (**Fig. 1C**). N form strongly affected the plant’s root and shoot fresh-weight biomass (**Fig. 1D**). The supply of +NO_3_^−^ resulted in the highest mean root fresh biomass at 137.96 mg, followed by +NH_4_NO_3_ treatment at 64.46 mg, while the +NH_4_^+^ supply resulted in the smallest root biomass at 35.39 mg. The +NH_4_NO_3_ supply resulted in the highest mean shoot biomass at 202.94 mg, followed by significantly lower +NO_3_^−^ treatment at 148.82 mg and +NH_4_^+^ treatment at 116.46 mg. The transfer to iN-conditions significantly increased the root:shoot fresh biomass ratio relative to the iN+ treatments by an average of 42 % for NO_3_^−^, 123 % for NH_4_NO_3_, and 158 % for NH_4_^+^ (**Fig. 1D**). The iN-resulted in significantly lower shoot N content at < 3.4 % than iN+ conditions at > 5.0 % (**Fig. 1E**). Shoot C content was statistically identical across iN treatments at means between 47.2 to 49 % (**Fig. 1F**). When comparing the iN+ treatments, the +NH_4_NO_3_ caused a statistically significant ^13^C enrichment of shoot *δ*^13^C values to −21.15 ‰ relative to the other two iN+ treatments at 21.81 ‰ for +NO_3_^−^ and −22.02 ‰ for +NH_4_^+^, while the iN-starvation did not significantly change shoot *δ*^13^C values relative to the iN+ counterparts (**Fig. 1G**). The shoot *δ*^15^N mean values differed between iN forms; the most ^15^N-enriched was +NH_4_NO_3_ at −1.83 ‰ compared to the other two iN+ treatments that were statistically lower at < −4.5 ‰ (**Fig. S2**). Additionally, the comparison between iN- and iN+ treatments showed statistical differences in *δ*^15^N values only for the NO_3_^−^ group (by an average of 1 ‰) but not for the other two treatment pairs (**Fig. S2**).

**Fig. 1.**
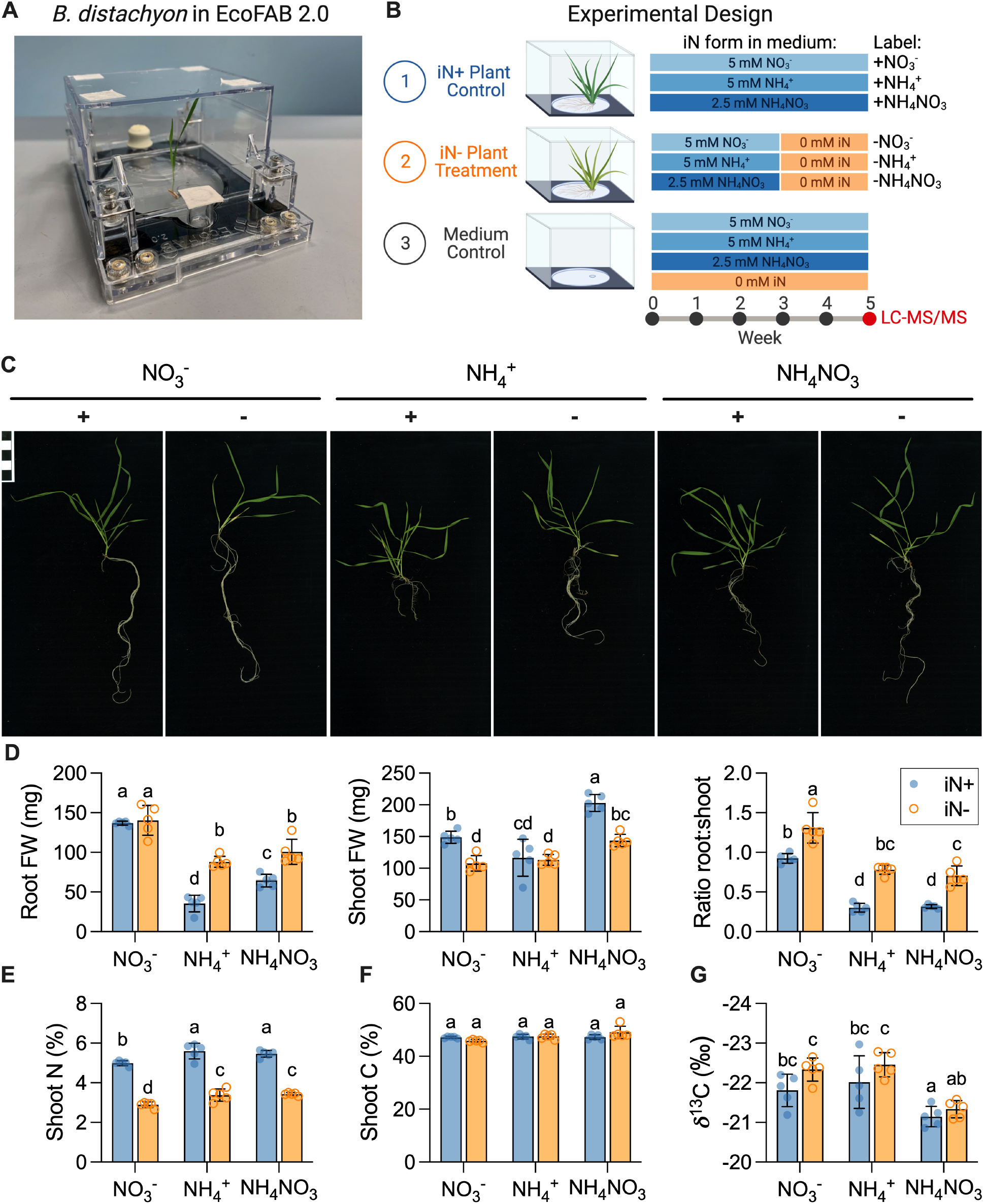
Analysis of plant biomass grown in EcoFAB 2.0. (**A**) An example of EcoFAB 2.0 containing hydroponically grown *Brachypodium distachyon* BD21-3 seedling. (**B**) Design of the plant growth experiment. *B. distachyon* Bd21-3 grew in EcoFAB 2.0 devices for 5 weeks with weekly media changes. Control iN+ plants (blue) received 5mM iN as nitrate, ammonium, or ammonium nitrate (labels: +NO_3_^−^, +NH_4_^−^, or NH_4_NO_3_), while the plants in the iN-treatment group (orange) were transferred to iN-medium (labels: −NO_3_^−^, −NH_4_^+^, −NH_4_NO_3_) after 3 weeks. The EcoFABs 2.0 filled with growth media without plants were used as technical controls. Each iN treatment had 5 biological replicates. We sampled plant spent media for exudate analysis by LC-MS/MS at week 5. The Fig. was created with BioRender.com. (**C**) The phenotypes of 5 weeks old *B. distachyon* grown on different iN forms. The iN+ controls are marked with plus signs (+), and the iN-treatments with minus signs (-). The scale bar in the upper left indicates 1×1 cm squares. All phenotype pictures are available at https://doi.org/10.6084/m9.figshare.21376017. (**D**) Fresh weight of roots, shoots, and their ratio. (**E**) Total nitrogen in dry shoot weight (% w/w). (**F**) Total carbon in dry shoot weight (% w/w). (**G**) Carbon stable isotope ratios of shoots. Bars show mean±SD. Different letters indicate statistically significant differences (two-way ANOVA with Tukey’s HSD test; *n* = 5; *p* ≤ 0.05). Blue-shaded circles show control iN+ plants, while orange circles indicate iN-plants.

### Inorganic nitrogen form modulates nitrogen uptake and rhizosphere pH

We measured the iN concentration in the spent plant growth media from the EcoFAB 2.0 root zones in the medium controls and plant treatments at the end of weeks 3 and 5. The NH_4_^+^ and NO_3_^−^ technical controls each contained, on average, 50 µmol of iN, which equals the intended 5 mM of iN. However, the NH_4_NO_3_ technical control contained, on average, 35 µmol of both NH_4_-N and NO_3_-N, totaling 7 mM of iN (40 % higher than the expected concentration of 5 mM). Plants did not take up all available iN from the growth media, leaving behind more than 36 % bioavailable iN each week (**Fig. 2A**). The iN plant uptake rates of solely supplied NO_3_^−^ or NH_4_^+^ were comparable at 25 or 28 µmol iN week^−1^ at week 3 and increased with plant age to 32 or 31 µmol iN week^−1^ at week 5, respectively. However, *B. distachyon* supplied with NH_4_NO_3_ preferred NH_4_^+^ over NO_3_^−^ in an approximate ratio of 2:1, uptaking on average 20 µmol NH_4_-N and 11 µmol NO_3_-N week^−1^ at week 3, and increasing to 25 µmol NH_4_-N and 12 µmol NO_3_-N week^−1^ at week 5 (**Fig. 2A**).

**Fig. 2.**
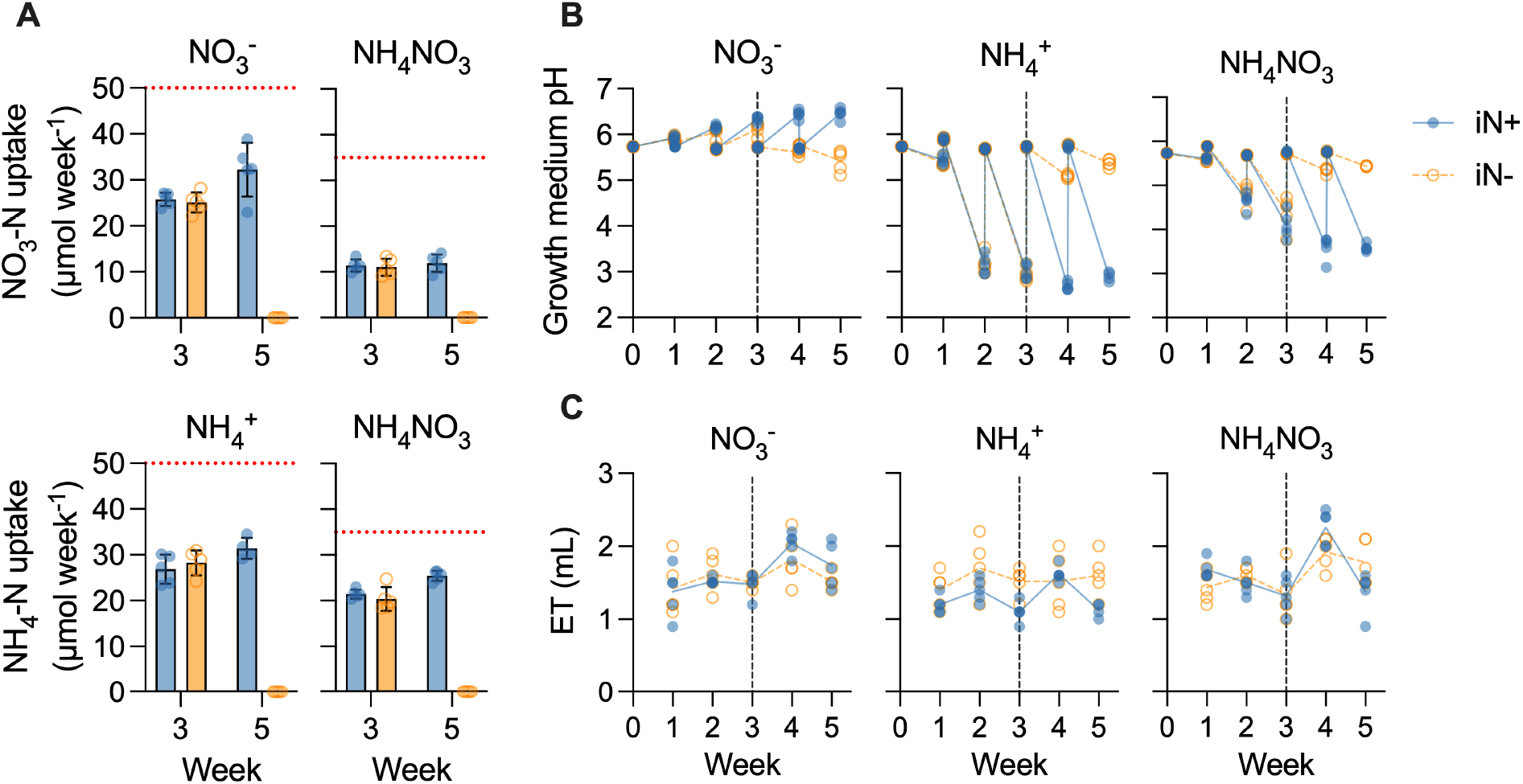
EcoFAB 2.0 growth medium characterization. (**A**) Plant nitrate-N and ammonium-N uptake (µmol/week) from the measured average concentration in the growth medium (dotted red line) for plants supplied with NO_3_^−^, NH_4_^+^, or NH_4_NO_3_. Bars show mean ±SD, *n*=5, +NH_4_^+^ treatment *n*=4. (**B**) Growth medium weekly pH shift from the average baseline value of 5.8. (**C**) Evapotranspiration (ET) expresses weekly media volume loss from EcoFAB 2.0 containing *B. distachyon*. The dashed vertical line at the end of week 3 indicates the transition to an iN-medium for the iN-treatment plants (empty orange circles and dashed lines), while the iN+ control plants received iN for 5 weeks (full blue circles and lines). The lines in panels b and c connect weekly means. Media (*V* = 10 ml) were changed completely weekly.

Each week, we measured the pH of the spent medium and replaced it with fresh medium pH of 5.8. Despite the presence of the MES buffer in the growth media, plants changed the media pH depending on the iN source, increasing the weekly pH amplitudes with age (**Fig. 2B**). The NO_3_^−^ uptake increased medium alkalinity to a pH of 6.5 at week 5, while NH_4_^+^ acidified the medium to a pH of 2.9 at week 5. NH_4_NO_3_ supply caused medium acidification to a pH of 3.6 at week 5 (**Fig. 2B**), correlated with a higher proportion of NH_4_^+^ to NO_3_^−^ uptake (**Fig. 2A**). Plants in the iN-free medium maintained the initial pH of 5.8 regardless of the previous iN supply (**Fig. 2B**). We have tested the growth of plants in MES buffer-free media, which resulted in quick acidification and severely reduced fitness of NH_4_^+^ supplied plants (data not shown).

Evapotranspiration was steady, averaging 1.6±0.2 mL week^−1^, i.e., 16% of medium initial volume (**Fig. 2C**).

### Root exudation changes with nitrogen supply

We analyzed the root exudate composition by LC-MS/MS (**Table S1**). The analysis was filtered down to 2065 features across positive and negative polarities, followed by the creation of molecular networks (**Fig. 3A, Table S2**). The network shows that the compositions of plant exudates are modulated by iN source with evident clusters of features differentially responding to iN supply (**Fig. 3A**). There were, in total, 155 features (7.8% of all features) with robust annotation, defined by MQScore (cosine score) > 0.7, to a library reference compound in GNPS across all samples/conditions (**Fig. 3A**). Many of these 155 features also had secondary GNPS annotations, most of which shared the same NPClassifier biosynthetic pathway as the top annotation (**Table S3**); this allowed for high confidence in accurate feature classifications. We show that 65 features were >1000-fold statistically higher (*t*-test, *p* ≤ 0.05) in the iN-(*n*=14) than in N+ (*n*=15) plant treatments (**Fig. 3A**). These top iN-features had no annotations in GNPS. Most top iN-features had relatively high mass-to-charge ratio (*m*/*z*) values >300. Notably, the top iN-features are clustered separately from the annotated features, thus prohibiting identification by mass shift (**Fig. 3A**). Because the highest *m*/*z* in our annotated features were flavonoids and carbohydrates (**Table S2**), we speculate that the top iN-unknowns with high *m*/*z* (**Fig. 3A**) may belong to these classes.

**Fig. 3.**
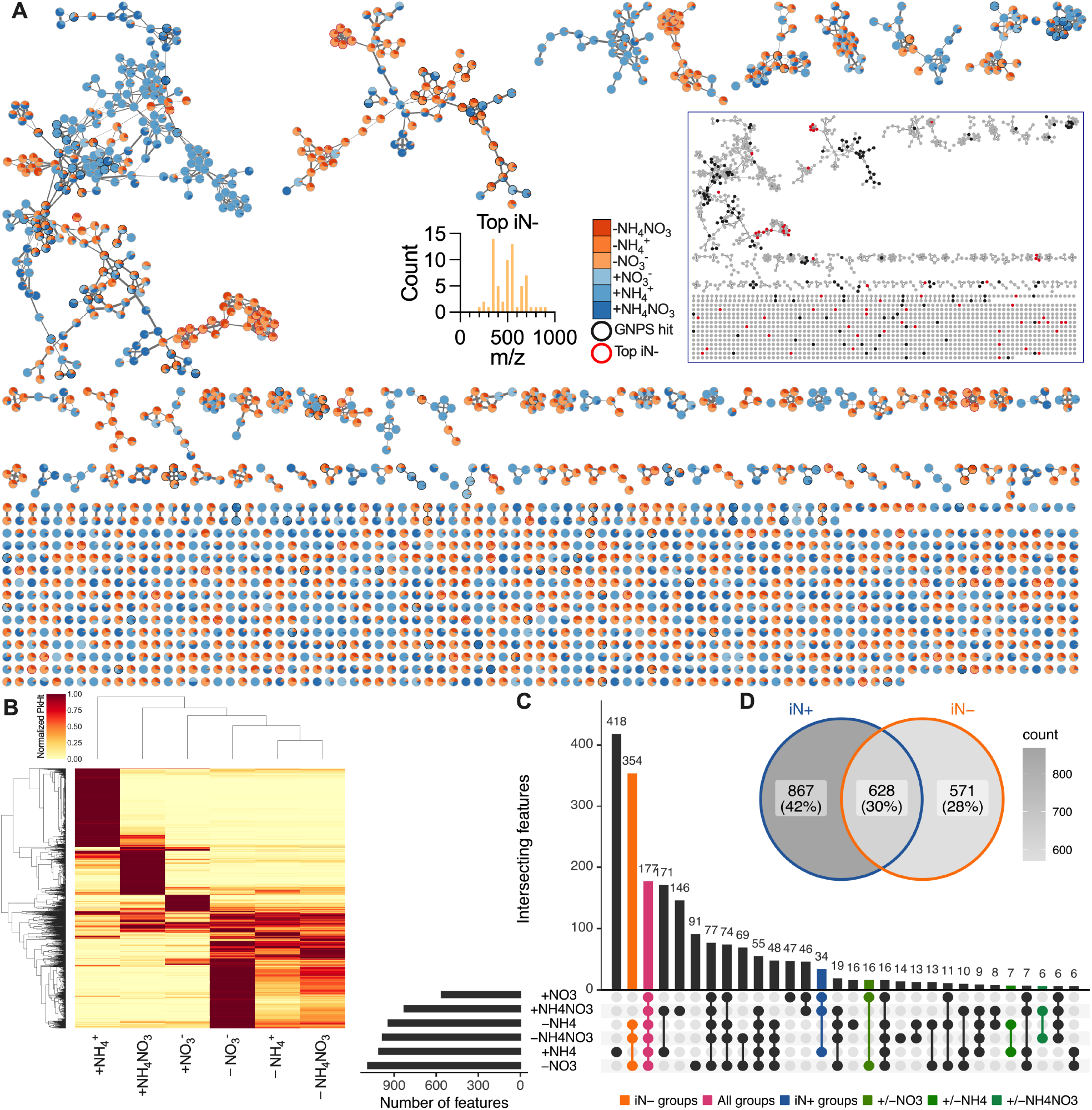
Filtered features detected by polar metabolomics in *B. distachyon* exudates under varying iN supply in EcoFAB 2.0. (**A**) Feature-based molecular networking (merged polarities, *n*=2065). Edge widths increase with the cosine score. Pie charts show the mean peak heights of iN+ (blue gradient) and iN-(orange gradient) treatments. The black border signifies GNPS annotated features (MQScore>0.7), while the red border shows 65 features that increased >1000-fold in N-deficiency (top iN-). Most top iN-features have *m*/*z* >300. Note no GNPS annotations for the top iN-features. The boxed network shows low proximity of the top iN-to GNPS features, thus prohibiting their mass-shift identification. The high-resolution interactive network is available at www.ndexbio.org under the UUID: 0e3c4384-35fa-11ed-ac45-0ac135e8bacf. (**B**) The heatmap shows hierarchical clustering of the filtered features normalized to the maximum average peak intensity across N treatments. The feature intensities across the iN+ treatments differed, while iN-formed a distinct cluster. (**C**) The UpSet plot shows the number of shared filtered features across the iN treatments among the first 30 largest intersecting sets. The second largest intersection of 358 features was across iN-treatments (orange), 193 features were shared across all iN groups (pink), 35 features were shared within the iN+ group (blue), while a relatively low number of features were shared between the individual iN+ treatments and their iN-counterparts (greens). (**D**) Venn diagram shows features intersections between grouped iN+ vs. iN-treatments.

We summarized the normalized relative feature peak height in a heat plot (**Fig. 3B**) where iN- and iN+ treatments clustered separately. Notably, the −NO_3_^−^ treatment showed high intensities for more features than −NH_4_^+^ or −NH_4_NO_3_ treatments. The feature intensities across iN+ treatments were different. In particular, +NH_4_^+^ and +NH_4_NO_3_ plant exudates contained more features with high intensities than the +NO_3_^−^ treatment. Additionally, the NMDS ordination plot showed separate clustering of iN- and iN+ treatments based on feature peak intensities (**Fig. S3**).

When we compared shared features across our 6 individual treatments, we observed minimal overlaps between the three iN+ treatments vs. their iN-counterparts (16 features for +NO_3_ vs. −NO_3_, 7 features for +NH_4_ vs. −NH_4_, and 6 features +NH_4_NO_3_ vs. −NH_4_NO_3_), while the three iN-treatments (−NO_3_ vs. −NH_4_ vs. −NH_4_NO_3_) shared a large proportion (354) of features (**Fig. 3C**). This suggests that the exposure of plants to iN-conditions cause reprogramming of exudation regardless of the previous iN form. The three iN+ treatments (+NO_3_, +NH_4_, and +NH_4_NO_3_) shared only 34 features, suggesting that different N forms cause different exudate profiles. The upset plot showed 177 features intersecting across all groups, thus potentially comprising core plant exometabolome. In addition, the number of features uniquely present only in one of the treatments (418 in +NH_4_^+^, 146 in +NH_4_NO_3_, 47 in +NO_3_^−^, 91 in −NO_3_^−^, 14 in −NH_4_NO_3_, and 16 in −NH_4_^+^) (**Fig. 3C**) corresponds with the highest peak intensity patterns in the heat plot (**Fig. 3B**).

The iN+ treatments collectively contained the highest number of features at 867 or 42%. The iN+ and iN-treatments shared 628 or 31% of all features, while 571 or 27 % were specific only to iN-condition (**Fig. 3D**). This suggests the highest cumulative feature diversity of polar metabolites under a sufficient iN supply.

The 155 annotated features belonged to diverse biosynthetic pathways according to the NPClassifer: amino acids and peptides (53), carbohydrates (39), alkaloids (27), shikimates and phenylpropanoids (22), fatty acids (8), no classification (4), terpenoid (1) and polyketides (1) (**Fig. S4**). These 155 annotations matched 110 unique metabolites (**Table S2**). There was a tendency (not statistically significant) for differential production of compound classes based on nitrogen supply (**Fig. 4A and B**). For example, signals for putatively annotated sugars were equally present across N treatments, and amino acid-annotated features were high under N-sufficiency. However, features annotated to Shikkimates/Phenylpropanoids had higher peak intensities in N-depleted conditions (**Fig. 4B**). Generally, under iN-conditions, the exudation of N-containing compounds was decreased relative to iN+ conditions, whereas the exudation of N-free compounds was increased relative to iN+ conditions (**Fig. 4B**). Specifically, iN-conditions increased the abundance of features with annotation to aromatic acids and their precursors (**Fig. S5**).

**Fig. 4:**
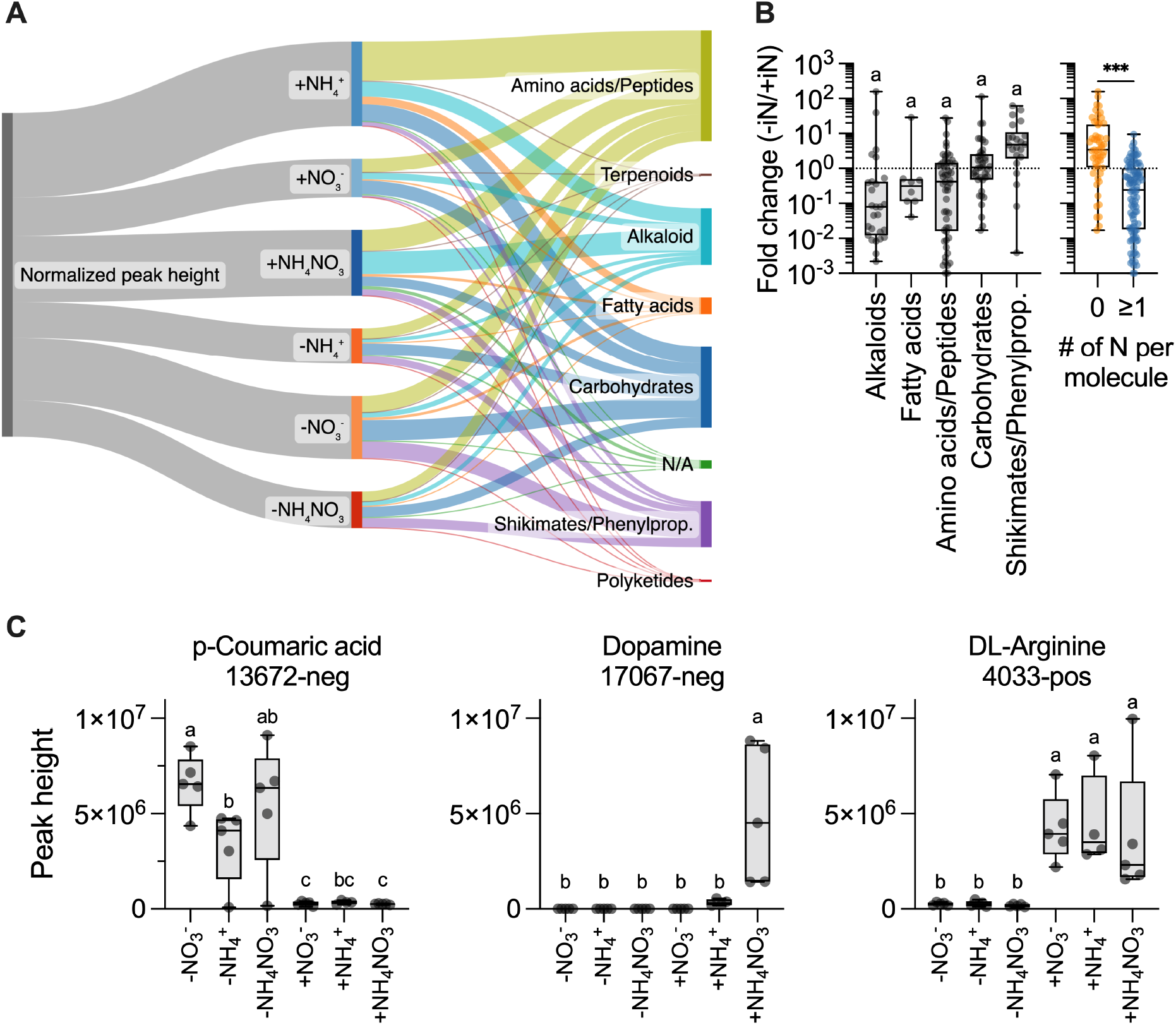
GNPS annotated features with MQScore > 0.7 (*n*=155) in the polar metabolomic analysis of *B. distachyon* exudates. (**A**) Sankey plot showing the distribution of compound classes for GNPS annotated features, which are classified by biosynthetic pathway and iN treatment. Widths represent the summed values of the normalized peak heights (relative to the max raw height value for each feature). The plot was made with SankeyMATIC (sankeymatic.com). (**B**) Fold-change (Fc) peak heights in iN-vs. iN+ treatments for annotated features grouped by biosynthetic pathways (excluded singleton and N/A groups). Different letters indicate statistically significant differences (ANOVA with post hoc Tukey’s HSD test, all n.s., *p* ≤ 0.05). There was a significant decrease in metabolites containing ≥ 1 N atom (blue) relative to N-free molecules (orange) under iN-conditions (t-test, *** *p* < 0.001). (**C**) Annotated features that showed differential patterns of abundance based on iN supply and were verified against library reference standards. The exudation of *p*-coumaric acid was increased in iN-, dopamine was +NH_4_NO_3_ specific, while DL-arginine increased only in iN+ conditions. Different letters above box plots indicate statistically significant differences (two-way ANOVA with post hoc Tukey’s HSD test; *n* = 5, +NH_4_^+^ *n* = 4, *p* ≤ 0.01). All box plots show all points, hinges extend from the 25th to 75th percentiles, the middle line indicates the median, and the whiskers extend to min and max values.

The supply of iN+ in the form of +NH_4_^+^ and +NH_4_NO_3_ resulted in the production of the largest number of unique features compared to +NO_3_^−^, whereas for iN-, the −NO_3_^−^ condition resulted in the largest number of features compared with −NH_4_^+^ and −NH_4_NO_3_ (**Fig. 3C**); these patterns were also present across the annotated features for relative normalized peak heights (**Fig. 4A**). In particular, the +NH_4_^+^ led to higher relative signals for Amino acids/peptides, while alkaloid biosynthesis was the highest under +NH_4_NO_3_ (**Fig. 4A**). Generally, the transition of plants from iN+ into iN-conditions caused a reduction in total normalized peak intensities, except for the +/−NO_3_^−^ treatments, however relative intensities of some molecular classes went up (shikimates in −NO_3_^−^ and +NH_4_NO_3_ and Carbohydrates in −NO_3_^−^ treatment).

Statistical comparison (ANOVA with post hoc Tukey’s HSD test, *p* ≤ 0.01) between peak heights of filtered features within iN-treatments, iN+ controls, and individual −/+iN pairs confirmed that exudation is modulated by iN source (**Table S4)**. The next aim was to verify some of the most interesting putative GNPS annotations. The statistical analysis combined with Upset grouping (**Fig. 3C**) of annotated features determined 6 features unique to iN-conditions, 2 in iN+, 10 specific to +NH_4_^+^, 4 in +NH_4_NO_3_, and 1 in +NO_3_^−^ (**Table S5**). These differentially produced features were matched against an in-house library of standard reference compounds analyzed using the same LC-MS/MS methods, resulting in 4 matches, of which 3 aligned to *m*/*z* (corresponding adduct), retention time, and MS/MS spectra of the reference (**Table S5**). Dopamine (feature 17067-neg) classified in the alkaloid pathway was uniquely present under the supply of +NH_4_NO_3_ (**Fig. 4C**). The *p*-coumaric acid (feature 13672-neg), classified as shikimate/phenylpropanoid, was significantly higher in iN-than iN+, while an opposing pattern was present for DL-Arginine (4033-pos), which significantly increased in iN+ conditions.

### Shikimic and *p*-coumaric acids affect the growth of bacteria and plants

Given the observed increase in features annotated as shikimates-phenylpropanoids under N-deficiency, we sought to assess their biological effects. We selected two representative compounds, *p*-coumaric and shikimic acid. Additionally, we used oxalic acid and glucose as controls because they are common in root exudates (*11*). The individual application of these four exogenous metabolites tested for effects on model rhizosphere bacteria *P. putida* KT2440 and *B. distachyon* Bd21-3 seedlings.

First, we investigated if the exogenous compounds can support improved bacterial growth. To see the effects of the additional C supply in the rich media, the bacteria grew for 72h. The OD_600_ of *P. putida* increased with increasing concentration (5, 50, or 500 µM) of shikimic acid in a liquid LB medium. However, oxalic acid, glucose, and *p*-coumaric acid had no bacterial growth-promoting effect (**Fig. 5A**).

**Fig. 5:**
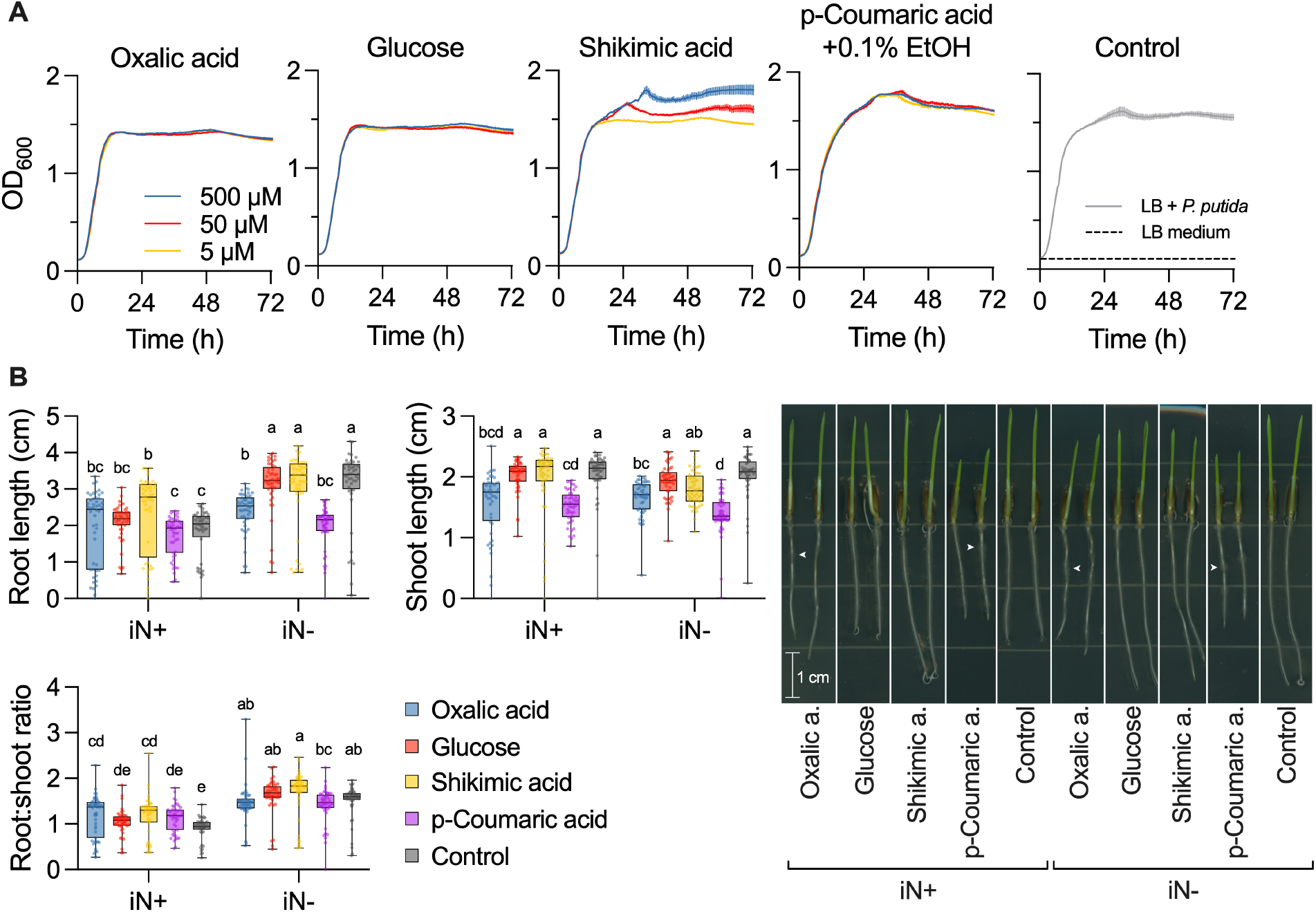
Exometabolite bioassays. Effects of exogenous application of root exudate compounds (oxalic acid, glucose, shikimic acid, *p*-coumaric acid) on (**A**) growth of *Pseudomonas putida* KT2440 (*c* = 5, 50, and 500 µM) in a multiwell plate and (**B**) 3 days old *Brachypodium distachyon* Bd21-3 seedlings (*c* = 500 µM) on phytoagar plates. (**A**) Shikimic acid increased microbial growth. (**B**) Shikimic acid increased root length under iN supply, while oxalic and *p*-coumaric acid decreased root length under iN starvation. The *p*-coumaric acid significantly decreased shoot length independent of iN supply. The iN was supplied at 2.5 mM of NH_4_NO_3_. Different letters indicate statistically significant differences (two-way ANOVA with post hoc Tukey’s HSD test; *n* = 48; *p* ≤ 0.05). The right panel shows representative seedlings’ phenotypes at day 3 since the onset of germination in the presence of the compounds. Arrows point to root areas with localized high root hair density in oxalic and *p*-coumaric acid treatments. All seedling pictures in high resolution are available at https://doi.org/10.6084/m9.figshare.21433065.

It is known that exometabolites can alter root growth (*6*); therefore, we also tested if these compounds at 0.5 mM affect seedling development under iN+ or iN-conditions (2.5 or 0 mM NH_4_NO_3_, respectively). The assay revealed that *B. distachyon* seedlings under the iN+ conditions statistically increased root length on shikimic acid to 2.36 cm relative to metabolite-free control at 1.80 cm. However, the iN+ condition did not follow this pattern (**Fig. 5B**). The *p*-coumaric acid supply significantly reduced mean shoot length relative to control from 1.99 cm to 1.51 cm (24 % reduction) in the iN+ condition and from 2.02 cm to 1.38 cm (32 % reduction) in iN-condition. However, the mean root length suppression by *p*-coumaric acid relative to the control was significant only in the iN-condition from 3.11 cm to 1.98 cm (37 % reduction) but insignificant in the iN+ condition (**Fig. 5B**). The oxalic acid biomass reduction relative to control followed the same pattern as *p*-coumaric acid by decreasing significantly shoot length in both iN+ and iN-conditions to 1.56 cm (22 % reduction) and 1.67 cm (18 % reduction), respectively; in addition to significant root length reduction to 2.38 cm (23 % lower) that was exclusive to iN-conditions. The oxalic and *p*-coumaric acids caused patchy root hair density by visual inspection (**Fig. 5B**).

### Reproducibility of *B. distachyon* measurements in EcoFAB 2.0 experiments

Several measurements were made during the experiment as described above, including: plant shoot and root biomass measurements, biomass elemental compositions, growth media NH_4_^+^ and NO_3_^−^ levels after 3 and 5 weeks of growth, pH and evapotranspiration, and untargeted LC-MS/MS feature comparisons. To assess reproducibility within this experiment, the coefficient of variation (CV; %) was calculated for each of these measurements under each experimental treatment and is detailed in (**Table S6**). In short, for 7/12 of the biomass measurements, the CVs were below 10%, all 12 were below 20%, and all N and C biomass composition measurements were below 10%. The flow injection analysis (FIA) measurements used to determine NH_4_^+^ and NO_3_^−^ concentrations after 3 and 5 weeks of growth had CV percentages almost exclusively below 20% except for the week 5 ‘−NO_3_^−^’ treatment with a CV of ~31%. All pH measurements, i.e., 6 conditions and 5 weeks, had CVs below 10%, and while evapotranspiration measurements varied more overall, none exceeded 30%.

As mentioned, after filtering based on intensity and statistical relevance, 2065 LC-MS features were used to compare *B. distachyon* exudate profiles across different nutrient growth media. Between 596 and 1159 of these features were detected under each experimental condition with an average peak height intensity greater than 5e5 (**Table S6**). Except for the ‘-NH_4_^+^’ condition, where 1 of the 5 replicates was removed, 22.6% to 31.2% of those features had a CV % of peak height below 30%.

## Discussion

### EcoFAB 2.0 is suitable for studying nitrogen nutrition impact on plant traits

We observed generally slower plant development manifested by the lack of flowering (**Fig. 1C**); this was due to the relatively short photoperiod of 12 h, which previously prohibited flowering in 35d-old *Brachypodium* Bd21-3 (*30, 31*). Keeping plants at the vegetative stage is advantageous due to EcoFAB’s compact size (**Fig. 1A**). Furthermore, the developmental stage is essential when comparing exudate data between studies; for example, the exudate profiles of another grass plant *Avena barbata* showed a shift with plant development (*3*).

The plants supplied exclusively with NH_4_^+^ developed typical symptoms of ammonium toxicity characterized by stunted growth (**Fig. 1C**) and the onset of leaf chlorosis and necrosis that occurred at week 3 (not shown). These ammonium toxicity symptoms have previously been observed for ammonium-grown *Brachypodium* (*16, 32*). Ammonium toxicity is caused by ionic imbalances, disturbance of pH gradients across cell membranes, or oxidative stress (*33*). Our observed alleviation of NH_4_^+^ toxicity by co-supply of NO_3_^−^ (**Fig. 1C and D**) can be explained by NO_3_^−^ ability to induce root signals that determine NH_4_^+^ tolerance, alongside alkalinization of rhizosphere during NO_3_^−^ uptake, which we also have observed (**Fig. 2B**) (*34*). In contrast, nitrate did not cause toxicity (**Fig. 1**), as excess is stored in plant vacuoles and used for biosynthesis when needed (*35*). The NO_3_^−^ fed plants developed bigger root biomass (**Fig. 1D**), perhaps via NO_3_^−^ driven regulation of auxin levels in roots (*17, 18*). However, more research must be completed to determine the causality between nitrate uptake, root development, and hormone homeostasis in *Brachypodium*.

Our iN-plants showed only minor chlorosis (**Fig. 1C**) combined with root foraging (**Fig. 1D**), suggesting mild N deficiency (*14*). For our plants, the shoot N content decreased in the N-free medium to <3% (**Fig. 1E**). According to data from closely related wheat, values <3.4% at the late tillering stage constitute N deficiency (*36*). Based on the reported critical N values in wheat and a general increase in root:shoot biomass under iN-conditions (**Fig. 1D**), our *Brachypodium* was N deficient. However, other studies observed lower N contents of ~1% for chronically undernourished (0.1 mM NO_3_^−^) 35-day-old *Brachypodium* (*15*). A previous study measured *Brachypodium* shoot N contents at the tillering stage at ~5% when supplied with sufficient NO_3_^−^ and ~6% for NH_4_^+^ or NH_4_NO_3_ (*16*). These values align with our results for N-supplied plants (**Fig. 1E**).

The high *δ*^13^C of +NH_4_NO_3_ plants (**Fig. 1F**) and comparable evapotranspiration across treatments (**Fig. 1D**) point to lower CO_2_ concentration inside the leaf or EcoFABs 2.0 chamber caused by higher C assimilation rates than in the other treatments. Our reasoning is based on the fact that carbon isotope ratios (^13^C/^12^C or *δ*^13^C) in C3 plants reflect leaf internal/atmospheric CO_2_ concentration, the long-term balance between CO_2_ supply and demand (*37*). The relationship between CO_2_ supply from the atmosphere versus plant demand is determined by stomatal conductance and assimilation rate, respectively. During CO_2_ fixation, kinetic isotope effects of RuBisCO result in preferential carboxylation of ^12^CO_2_. If the leaf’s internal CO_2_ concentration drops, the ^13^CO_2_ is increasingly fixed, resulting in an increase of leaf *δ*^13^C (*37*). We can assume high stomatal conductance across the treatments at high water supply and relative humidity in EcoFAB (>90% at week 5, data not shown).

### Nitrogen form determines rhizosphere pH that can influence root exudation

The higher uptake of NH_4_ relative to NO_3_ for *Brachypodium* (**Fig. 2A**) was shown previously (*38*). The iN form drove bidirectional pH change in our spent medium (**Fig. 2B**). Plant uptake of ammonium decreases pH in the rhizosphere, while nitrate causes an increase in pH due to their root-mediated efflux of H^+^ or OH^−^, respectively (*39*). We could not separate the pH and iN form effects in our experiments. However, this iN form-associated pH change is a natural process. Plants can alter the pH in a sharp gradient from the root surface (2–3 mm), and the size of this layer increases with lower soil buffering capacity. This interface is a hotspot for exudation and plant-microbe interactions (*40*). Our observations are, therefore, relevant from the perspective of natural rhizosphere processes. The acidification during NH_4_^+^ and NH_4_NO_3_ treatments (**Fig. 2B**) might cause a high number of unique exudate features (**Fig. 3C**) in comparison to alkalinized rhizosphere under NO_3_^−^ supply (**Fig. 2B**). It is well-established that proton gradients across the membranes facilitate the transport of compounds into and from the rhizosphere (*41*). For example, the exudation of terpenoid 3-epi-brachialactone in grass *Brachiaria humidicola* was promoted during low rhizosphere pH and NH_4_^+^ nutrition due to changes in proton motive force (*42*).

### Nitrogen sufficiency and deficiency in grasses produce conserved patterns of root exudation

Metabolomic analyses revealed dramatic changes in exudate composition in response to nitrogen supply (**Fig. 3, 4, and S5**). Many of the detected top annotations (**Fig. S4, Table S2**) align with organic, aromatic, or amino acids previously reported in root exudates of plants (*3, 43, 44*). Therefore, suggesting the relevance of the annotations.

*B. distachyon* supplied with iN+ increased abundance of N-containing compounds, which is exemplified by increased alkaloids and amino acid annotations; this was in contrast to iN-treatments that decreased these compounds (**Fig. 4A and B**). Similarly, switchgrass showed an increased amino acid exudation pattern under +N and +NP fertilization relative to unfertilized controls (*20*). Additionally, axenically grown maize under N deficiency lowered the amino acid amounts in root exudates (*22*).

*B. distachyon* produced dopamine in the root exudates in NH_4_NO_3_ treatments (**Fig. 4C**). The number of studies showing dopamine production in plant tissues is limited (*45*). Dopamine has a range of physiological and biochemical functions in plants, including tolerance against abiotic stresses, such as drought, salt, and nutrient stress (*45*). A recent study showed that exogenous dopamine could mediate nitrogen uptake and metabolism regulation at low ammonium levels in a tree *Malus hupehensis* (*46*). To our knowledge, dopamine exudation in grasses remains to be shown experimentally. However, the production of another neurotransmitter, serotonin, was documented to be also N-dependent in the rhizosphere of switchgrass (*20*).

We have observed a fold increase in iN-treatments for the shikimate-phenylpropanoid pathway metabolites (**Fig. 4 and S5**). According to NPClassifier, the shikimate/phenylpropanoid pathway contains shikimic acid precursors (e.g., quinic acid) and derivatives, aromatic acids (e.g., cinnamic acid derivatives), coumarins, hydroxybenzoic acids, flavonoids, lignans, or lignins (*27, 47*). Keeping this classification in mind, the literature contains several examples where shikimates-phenylpropanoids increased in the exudates of grasses under N deficiency. Recent work showed an increase in several organic and aromatic acids in marginal N-limited soil of the switchgrass rhizosphere (*20*). Another paper showed that the root exudation of *p*-coumaric acid in rice increased by 1.95-fold in iN-than iN+ conditions (*21*). Therefore, our results from *B. distachyon* combined with the previous result suggest conserved patterns of increased production of compounds from the shikimate-phenylpropanoid pathway in root exudates of grasses under N-deficiency (*20, 21*).

### Shikimic versus aromatic acids have diverse functions in the rhizosphere

We observed *in vitro* growth-promoting effects of shikimic acid on *P. putida* KT2440 (**Fig. 5A**). This is consistent with other studies showing that rhizosphere bacteria prefer to consume shikimic acid alongside other organic acids (nicotinic, salicylic, cinnamic, and indole-3-acetic) exuded by roots of grass *Avena barbata* (*3*). Moreover, shikimic acid was shown to be a highly consumed substrate by diverse phylogeny of soil isolates in the recently developed Northen Lab Defined Medium (NLDM) (*48*). The shikimic acid promoted the growth of the roots only under iN+ conditions relative to a control (**Fig. 5B**). The root growth-promoting effects of 0.5 mM shikimic acid in the presence of 0.5 mM NH_4_NO_3_ were also shown in 10 days old *Arabidopsis* seedlings (*6*). Synthesis of previous observations and our exometabolomics, microbial, and root growth data (**Fig. 3, 4, and 5**) suggest that exogenous shikimic acid can promote root growth, serve as substrates to support the growth of soil bacteria, and thus shape microbiome assembly in the rhizosphere (*3, 6, 20, 48*).

We found no effect of increasing the concentration of *p*-coumaric acid on *P. putida* growth (**Fig. 5A)**. This is consistent with a recent study that showed that coumaric acid is generally not a preferred microbial substrate by soil bacterial isolates *in vitro* (*49*). Our data suggest that concentrations of 500 µM and higher can have phytotoxic activity on *Brachypodium* seedlings and may negatively impact crop productivity (**Fig. 5B**). In particular, *p*-coumaric acid strongly inhibited root growth exclusively under iN limitation (**Fig. 5B**). It was shown that N-limiting conditions induce root foraging via the regulation of auxin transport (*14, 17, 18*). Therefore, we propose that *p*-coumaric acid in *Brachypodium* might alter root hormone homeostasis, likely via stimulation of IAA oxidase activity shown for other aromatic compounds (*5, 50*). In addition, the inhibition of shoot growth was independent of N supply (**Fig. 5B**). The shoot reduction might be related to photosynthesis alterations via peroxidase-mediated chlorophyll degradation shown for *p*-coumaric acid previously (*51*).

### Limitations and future work

The current experimental design compromises sufficient growth conditions and exudate signals. In our setup (EcoFAB 2.0), some compounds might not accumulate in detectable levels in our exudate samples due to the need for weekly medium changes to maintain adequate N supply and acceptable pH (**Fig. 2**). Growing *B. distachyon* in a larger medium volume than 10 ml to avoid pH shifts would dilute the exudates concentration. Moreover, metabolite extraction from the greater volume of spent media would result in higher ion suppression due to high salts and MES buffer concentrations. While EcoFAB 2.0 can have an opaque base to shield the roots, this study used a transparent base; thus, the light exposure could have impacted the exudate profiles due to stress. The compact EcoFAB size limits the number of plant species to small plants. Grasses larger than *B. distachyon*, like wheat, rice, maize, sorghum, or switchgrass can be grown only during the early seedling developmental stages to avoid crowding inside the EcoFAB 2.0 device. The future EcoFAB designs will aim at accommodating larger plant species to the later developmental stages.

As mentioned above, the interconnected effects of the N source and rhizosphere pH (**Fig. 2**) have combined effects on the exudate composition (**Fig. 3**). From this perspective, it would be interesting to assess the sole effect of pH gradient on exudate composition in plants grown with one nitrogen source (i.e., NH_4_NO_3_). A controlled study establishing causality between pH and root exudate composition still needs to be conducted in grasses. Furthermore, future work elucidating dopamine pathways and exudation in grasses would benefit our understanding of grassland function under N-limitation and stress.

The typical GNPS annotation rate is 5-6% (*52*). Thus unsurprisingly, the top fold-increasing iN-features were unannotated (**Fig. 3C**). We can not reveal the identity of these “dark matter” molecules potentially due to the lack of reference spectra in library databases (*53*). The future expansion of reference libraries will significantly increase the GNPS annotation rates.

We conclude that the new EcoFAB 2.0 devices result in reproducible plant growth, which enabled us to determine that the iN form (NO_3_^−^, NH_4_^+^, or NH_4_NO_3_) and starvation are significant factors modulating root exudation in the grass species *Brachypodium distachyon*. Relative to the +NH_4_^+^ and +NO_3_^−^ treatments, the +NH_4_NO_3_ supply produces large shoots, robust root exudate profiles (signal intensity and feature diversity), including dopamine, increased photosynthetic rate, balanced rhizosphere pH changes, and preferential uptake of NH_4_^+^ relative to NO_3_^−^. Transient iN-limiting conditions for two weeks re-modulate root exudation relative to the previous iN forms. The root exudate profiles of plants supplied with NO_3_^−^, NH_4_^+^, or NH_4_NO_3_ differed from each other and were collectively more diverse than the exudates of the iN-starved plants. The N-deficient plants increased the abundance of compounds in the shikimic-phenylpropanoid pathway, e.g., *p*-coumaric acid, while decreasing the abundance of N-containing compounds. We demonstrate on the example of shikimic and *p*-coumaric acids that the exometabolites increasing in the rhizosphere of N-stressed plants can serve diverse functions from microbial substrates to allelochemicals. Our work increases the understanding of nitrogen nutrition and root exudation in the model plant *B. distachyon*, which can be useful for studies of economically important grass crops. Furthermore, EcoFAB 2.0 device proved to be a valuable standardized tool in plant research and has the potential to enable scientists to work on shared systems and facilitate replicability in environmental microbiome research.

## Materials and Methods

### Plant growth and harvest

Seeds of *Brachypodium distachyon* Bd21-3 with removed lemma were surface-sterilized with 70% (v/v) ethanol for 30 s followed by 5 min in 6% (w/v) sodium hypochlorite and washed 5 times with milli-Q water. Seeds were then stratified on plates composed of 1.5% (w/v) Phytagel™ and incubated at 4°C for 3 days in the dark. Germination was done by placing plates vertically in a growth chamber (CU36L4, Percival Scientific Inc, Perry, IA, USA) set to 22°C, photosynthetic photon flux density (PPFD) at 150 µmol m^−2^ s^−1^ and a 12 h photoperiod. We transferred 3-day-old seedlings aseptically into novel fabricated ecosystem (EcoFAB 2.0) devices (**Fig. 1A**). EcoFAB 2.0 devices were assembled and sterilized following the instructions described in the supporting information (**Protocol S1**).

We supplied each EcoFABs with 10 mL of plant growth media modified from Glazovska *et al*.*(16)*. The basal salt medium (BSM) consisted of 0.06 µM MoNa_2_O_4_, 0.3 µM CuSO_4_, 0.8 µM ZnSO_4_, 9 µM MnCl_2_, 50 µM H_3_BO_4_, 50 uM Fe(III)-EDTA-Na, 0.5 Mm KH_2_PO_4_, 1mM MgSO_4_, 5mM MES buffer at pH 6. From the BSM, we created 4 types of media with varying iN composition: (i) the NO_3_^−^ medium contained 2mM Ca(NO_3_)_2_ and 1mM KNO_3_; (ii) the NH_4_^+^ medium contained 2.5 mM (NH_4_)_2_SO_4_; (iii) the NH_4_^+^/NO_3_^−^ medium contained 2.5 mM NH_4_NO_3_; and (iv) iN-medium consisted of BSM without additional ingredients. All iN-replete media contained iN at 5 mM. All media were filtered via 0.22 µm PES membrane and stored at 4 °C.

The iN-treatment, iN+ plant control, and medium control received their respective iN media (**Fig. 1B**). The EcoFABs 2.0 were placed into the growth chamber, continuing the settings during germination. We used 5 biological replicates resulting in a total of 50 EcoFABs 2.0. The plant growth media was entirely changed weekly in a sterile environment of a biosafety cabinet by opening EcoFAB sampling ports and using 10 ml sterile serological pipettes. Treatment group plants received iN+ medium for 3 weeks before being transferred to iN-medium for 2 weeks. We checked the sterility of all EcoFABs 2.0 by incubating a drop of spent media from weeks 3 and 5 on LB agar plates for 3 days at 30°C.

Plant harvest occurred at week 5 by dismantling EcoFAB 2.0 and gently pulling plants out of the root chamber opening. Autoclaved wipers (Texwipe® TX604 TechniCloth®) removed residual medium from the roots, and the whole plants were scanned (HP Scanjet G4050). The determination of *Brachypodium* developmental stages followed the work by Hong et al. 2011 (*30*). The roots and shoots were separated, and fresh biomass was immediately weighed. Roots were flash-frozen in liquid N_2_, while shoots were frozen by placing them at −80 °C. Frozen tissues were stored at −80 °C. Lyophilization (Labconoco FreeZone 2.5) removed water from the frozen shoots before homogenization using a beat mill (MM 400, Retsch GmbH, Haan, Germany) set to 30 Hz for 10 min.

### Spent medium sampling and analyses of pH and iN

We sampled the spent medium from EcoFAB 2.0 weekly and quantified evapotranspiration (ET) as a medium volume loss. We adjusted the medium samples to the initial 10 mL with milli-Q water and then sterile-filtered using Acrodisc® Syringe Filters 25 mm with 0.2 µm Supor® PES Membrane (Pall Corporation, NY, USA) attached to a 10 ml plastic syringe. We split each medium sample into 3 aliquots: 1 mL for pH measurement, 1 mL for flow injection analysis (FIA), and 8 mL for LC-MS. The pH was measured weekly using HALO® Wireless pH Meter (HI10832, Hanna Instruments). The FIA was done on the samples from weeks 3 and 5, while LC-MS/MS was done on samples from week 5. One of the plant control +NH_4_^+^ medium samples (EcoFAB #22) at week 5 was lost during sampling; thus, it was excluded from the downstream analyses resulting in *n*=4 for this treatment.

The analytical Facility at UC Davis conducted the FIA analysis. The method involved the quantification of ammonium (NH_4_-N), nitrate (NO_3_-N), and nitrite (NO_2_-N) in aqueous samples. The spent medium sample was diluted 10x from 1 mL to 10 mL with milli-Q water and used for the FIA (Lachat Quikchem 8500 Series 2). The detection limit for the 10x dilution was 0.5 mg/L for all analytes. The reported nitrate values include nitrite because nitrite is typically an insignificant fraction of the N in a sample. We calculated plant inorganic nitrogen uptake (*N*_uptake_) (µmol) (**Fig. 2A**) from 10 mL of the growth medium (*V*) using the measured initial iN concentration (mg/L) in the fresh medium (*c*_i_) and the final concentration in spent medium (*c*_f_) and molar mass of nitrogen (*M*_N_ = 14.0067 g/mol) according to **Equation 1**:

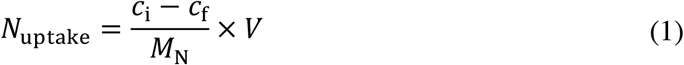

### Analysis of shoot elemental and stable isotope compositions

We conducted shoot C and N elemental (% dry weight) and isotopic composition (*δ*^13^C and *δ*^15^N values, respectively) analysis at the Center for Stable Isotope Biogeochemistry, University of California, Berkeley, CA, USA. The analytical system consisted of a CHNOS elemental analyzer (EA) (vario ISOTOPE cube, Elementar, Hanau, Germany) coupled to a continuous flow stable isotope mass spectrometer (IRMS) (IsoPrime 100 mass spectrometer, Isoprime Ltd, Cheadle, UK). The dried powdered shoots were packed into tin foils and analyzed. We report the stable isotope abundances as delta (*δ*) notation in parts per thousand (‰) according to **Equation 2**:

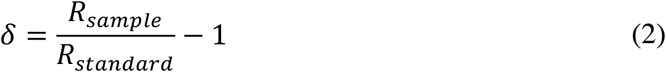

Where *R*_sample_ and *R*_standard_ are the ratios of the heavy to light isotope (i.e., ^15^N/^14^N, or ^13^C/^12^C) in the sample versus an international standard, respectively. The international standards are atmospheric nitrogen (air) for *δ*^15^N values and Vienna PeeDee Belemnite (VPDB) for *δ*^13^C values. Raw stable isotope data were corrected for drift and linearity and normalized to the international stable isotope reference scale. The normalization used three laboratory reference materials with different carbon and nitrogen delta values. These laboratory reference materials are calibrated annually against IAEA (International Atomic Energy Agency, Vienna, Austria) certified reference materials. We used NIST (National Institute of Standards and Technology, Gaithersburg, MD, USA) SMR 1577c (bovine liver) previously calibrated against IAEA-certified reference materials for quality control. The long-term external precision for C and N isotope analyses was ± 0.1‰ and ± 0.2‰, respectively.

### LC-MS analysis of *Brachypodium* exudates

The root exudate samples from week 5 (8 mL) were frozen at −80°C and then freeze-dried (Labconco Freeze-Zone). The dried material was collected by rinsing tubes in 700, 500, and then 300 uL of LC-MS grade methanol (Sigma); rinse volumes were combined in 2mL tubes and dried in a Thermo Speed-Vac concentrator. We resuspended the dried material in 150 uL LC-MS grade methanol containing internal standards (**Table S1**). The solution was vortexed 2×10 seconds, bath sonicated in ice water for 15 min, centrifuged (10,000 x g, 5 min, 10°C) to pellet insoluble material, and then supernatants were filtered using 0.22 um PVDF microcentrifuge filtration devices (Pall) (10,000 x g, 5 min, 10°C); filtrates were transferred to 384 well plates, and sealed with foil prior to analysis. Metabolites were separated using hydrophilic interaction chromatography for polar metabolomics; eluted compounds were detected using a Thermo Q Exactive Hybrid Quadrupole-Orbitrap Mass Spectrometer. LC-MS/MS parameters are in **Table S1**.

### Untargeted metabolomics and molecular networking

The GNPS and MZmine2 version 2.39 workflow generated molecular networking files for positive (12109 features) (MZmine2 parameters in **File S1**) and negative (4770 features) (MZmine2 parameters in **File S2**) polarities (*26*). First, a baseline filter accepted features with RT>0.6 min (post solvent front) and maximum peak height >1e6 and >10x the peak height maximum in extraction (*n*=10) and technical controls (*n*=5). We added +1 to the numerator and denominator in the fold-change calculations to avoid errors when dividing by 0. Additionally, we applied a statistical filter (ANOVA) accepting only features significantly higher (*p* ≤ 0.01) in plant samples than extraction and technical controls. Using Cytoscape software (*54*), we conducted feature-based molecular networking by merging the filtered features from positive and negative polarities into a single molecular network (2065 nodes) and following a step-by-step protocol (*52*). We selected the top iN-features based on the statistically significant and >1000 fold-increase in iN-(*n*=14) versus iN+ (*n*=15) treatments in exudates at week 5 (*p* ≤ 0.05, t-test). We accepted top GNPS annotations with cosine scores of MS/MS spectral mirror match (MQScore) to library reference >0.7 (155 features) (**Table S3**). NPClassifier determined metabolite classifications based on annotated features’ biosynthetic pathways (*27*). For each feature, we calculated the fold change (Fc) of average peak height in iN-treatments vs. iN+ in exudates at week 5. We determined in which treatment groups (6 treatment groups = UpSet, or 2 treatment groups = Venn) the filtered features were present based on their max peak height in the group exceeding a threshold of 1e6 (**Table S2**). For filtered features, we compared the peak height between plant treatments for statistical differences (ANOVA with post hoc Tukey’s HSD test, *p* ≤ 0.01 using the *pairwise_tukeyhsd* function from the statsmodels (v. 0.11.1) python package) (**Table S4**). For annotated features that showed differential abundances between iN treatments (based on upset grouping and statistical differences), we verified the feature’s top annotation against library reference in the metabolite atlas by matching RT (max Δ0.6 min), adduct, *m*/*z* integer, and MS/MS spectra of the reference (**Table S5**) (*55*).

### Growth of bacteria on selected exudate compounds

We selected representative compounds from shikimate (shikimic acid) and phenylpropanoid (*p*-coumaric acid) pathways to test their impact on the growth of soil bacterium *Pseudomonas putida* KT2440 (ATCC 47054), which was shown to colonize plant roots (*56*). We resuspended the bacteria to an OD_600_ (1 cm pathlength) of 0.01 in 2.5% LB (Luria-Bertani) liquid medium supplemented with a phosphate buffer (6g/L Na_2_HPO_4_, 3g/L KH_2_PO_4_) pH 7 and containing either oxalic acid, glucose, shikimic acid, or *p*-coumaric acid at 5, 50, or 500 uM (*n*=7). The *p*-coumaric acid treatments also contained 0.1 % ethanol, which was used as a solvent. Our controls were either the blank medium or the medium with the bacteria (*n*=6). A 96-well microtiter plate-based method using a Synergy H1 (BioTech Instruments) reader was used to monitor bacterial growth at 28°C with shaking through periodic measurements of changes in OD at 600 nm for 72 h.

### Growth of *B. distachyon* seedlings on selected exudate compounds

Dehusked and surface-sterilized seeds of *B. distachyon* BD21-3 were placed on 10 cm square plates containing N-free ½ MS salts in 0.75% (w/v) phytoagar containing no metabolite (control) or 0.5 mM oxalic acid, glucose, shikimic or *p*-coumaric acid and either 0 or 2.5 mM NH_4_NO_3_. The *p*-coumaric acid treatments also contained 0.1 % ethanol as a solvent. Each treatment was done in 4 replicate plates, each containing 12 seeds; this resulted in a total of 48 seedlings/treatment. The plates with seeds were stratified for 3 days at 4°C in the dark and then germinated vertically for 3 days in the growth chamber using the previous *Brachypodium* growth settings listed in the previous section. The root and shoot length for each plant were analyzed using ImageJ 1.53k software by setting the scale of each image to 1 cm and using the segmented line function to trace the center of the tissue from the embryo pole to the tip of the primary root and from the embryo pole to the tip of the first leaf, respectively.

### Statistical analyses and data visualization

We conducted feature filtering and t-tests between iN-vs. iN+ treatments in Microsoft Excel version 16.66.1. Data visualization and statistics were done in Prism 9 (GraphPad Software LLC) version 9.3.0 unless indicated otherwise. We checked the data for normality and followed ANOVA with post hoc Tukey’s HSD test when performing ANOVA analyses. Rstudio generated UpSet and non-metric Multi-dimensional Scaling (NMDS) plots using the UpSetR, and vegan packages, respectively (*57, 58*). The heatmap was generated using the Seaborn package (0.10.1) in Python (3.8.3). The statistical analysis of peak height data across plant treatments was also done in python using ANOVA with post hoc Tukey’s HSD test using the ‘pairwise_tukeyhsd’ function within Statsmodels (0.11.1) (**Table S4**). The Sankey plot was generated with the online tool SankeyMATIC (sankeymatic.com). The statistically significant difference was assumed at *p* ≤ 0.05 except for ANOVA of peak height data, which was performed at a threshold of *p* ≤ 0.01 following false discovery rate correction (*59*). To compare the variation of quantitative traits measured in this study, we used the coefficient of variation (CV) which was calculated as the standard deviation of replicate samples divided by their mean and then converted to percentage (*60*).

## Supporting information

Protocol S1

## Acknowledgments

The Analytical Laboratory at UC Davis conducted the FIA analysis. We thank Todd Dawson and Stefania Mambelli from the Center for Stable Isotope Biogeochemistry at UC Berkeley for elemental and stable isotope analysis. Silvia Glazovska from the Department of Plant and Environmental Sciences at the University of Copenhagen provided excellent advice for hydroponic media formulation.

## Author contributions

VN designed the research with input from PFA, TRN, KZ, BPB, and SMK. PFA developed the EcoFAB 2.0 device and assembly protocol. VN, PFA, YD, and ANG performed the research. VN, PFA, BPB, TRN, and SMK analyzed, collected, or interpreted data. VN wrote the first draft of the manuscript with input from PFA, CT, and TRN. All authors (VN, PFA, BPB, YD, KZ, CT, ANG, SMK, and TRN) edited the paper.

## Data and materials availability

GNPS negative and positive modes for the polar metabolite analysis (HILIC) are available at https://gnps.ucsd.edu/ProteoSAFe/status.jsp?task=7e8210b68fd049198579c2eff3068876 and https://gnps.ucsd.edu/ProteoSAFe/status.jsp?task=15c14ac5a13c466db52068de85bc8894. The interactive feature-based molecular network and node metadata can be accessed via Network Data Exchange (NDEx) at www.ndexbio.org under the UUID: 0e3c4384-35fa-11ed-ac45-0ac135e8bacf. Phenotypes of plants grown under varying iN supplies are available at https://doi.org/10.6084/m9.figshare.21376017. Phenotypes of seedlings grown on selected root exudate compounds are available at https://doi.org/10.6084/m9.figshare.21433065. All raw data used in this paper are available at https://doi.org/10.6084/m9.figshare.21440748.

## Supplementary Materials

**Fig. S1: EcoFAB 2.0 devices can accommodate a wide diversity of plant research**. (**A**) Empty EcoFAB 2.0 and its main functional parts. (**B**) Configurations of EcoFAB 2.0 for *B. distachyon* growth include (from left): EcoFAB 2.0 filled with liquid medium for hydroponic plant growth (used in this study), soil-filled setup, and a version of EcoFAB 2.0 with opaque black root chamber that allows light-sensitive rhizosphere experiments. (**C**) Examples of possible applications include (from left): gas analysis, microscopy of roots (2X and 40X magnification), root system visualization, study of microbiomes by 16S sequencing, and metabolomics on growth medium (used in this study).

**Fig. S2: Shoots N stable isotope ratios**. Bars show mean ±SD of iN+ of control (blue) and iN-(orange) plants. Different letters indicate statistically significant differences (two-way ANOVA with post hoc Tukey’s HSD test; *n* = 5; *p* ≤ 0.05).

**Fig. S3: NMDS plot for *B. distachyon* root exudate features using raw peak height data (filtered features, *n* = 2065)**. Each biological replicate across iN treatments is represented with a symbol. The Blue symbols represent iN+ treatments, while the orange symbols represent plants grown in iN-medium.

**Fig. S4: Sub-network of grouped annotated features**. The features were grouped into biosynthetic pathways by NPClassifier. Pie charts show relative mean peak heights for individual iN-treatments (shades of orange) and iN+ treatments (shades of blue). The network shows merged features from positive and negative polarities. The black borders indicate GNPS annotation with MQScore > 0.7 (*n*=155). The features represent results of polar metabolite analysis in root exudates of *B. distachyon* Bd21-3 at week 5.

**Fig. S5: Sub-network of annotated features abundant in N-deficient root exudates**. The network shows features significantly increasing > 5-fold in iN-treatments (*n*=15) relative to the iN+ treatments (*n*=14) (t-test, *p*-value ≤ 0.5). Pie charts show relative mean peak heights for individual iN-treatments (shades of orange) and iN+ treatments (shades of blue). The black borders indicate GNPS annotation with MQScore > 0.7. The network shows merged features from positive and negative polarities of polar metabolite analysis in root exudates of *B. distachyon* Bd21-3 at week 5.

**Protocol S1: Assembly and Sterilization of EcoFAB 2.0 for plant growth experiments.**

**Table S1: LCMS parameters and Metabolite extraction internal standards mix**.

**Table S2: Molecular network metadata node table.**

**Table S3: All GNPS annotations for features with top annotation MQ score > 0.7**.

**Table S4: Statistically significant differences (ANOVA with post hoc Tukey HSD test, p ≤ 0.01) between peak heights of features in plant exudate samples**.

**Table S5: Selection of differentially produced features and verification against library references**.

**Table S6: Coefficient of variation (CV) summary of all data**.

**File S1: Positive mode MZMine2 parameters.**

**File S2: Negative mode MZMine2 parameters**.

