## Supplementary material for "Reproducible growth of *Brachypodium distachyon* in fabricated ecosystems (EcoFAB 2.0) reveals that nitrogen form and starvation modulate root exudation": Protocol S1

| | |
|---|---|
| Protocol name | Assembly and Sterilization of EcoFAB 2.0 for plant growth experiments |
| Version | 2 |
| Date last updated | 20220927 |
| Editor(s) | Peter Andeer |
| Brief changes | Updated and formalized using current parts |
| Effective date | 20220927 |

### Introduction, intended use and scope

The following instructions are for the sterilization, and assembly of EcoFAB 2.0 for studying small plants and their microbiomes during growth.

### Materials

1. *EcoFAB 2.0 parts supplied (quantity per device)*

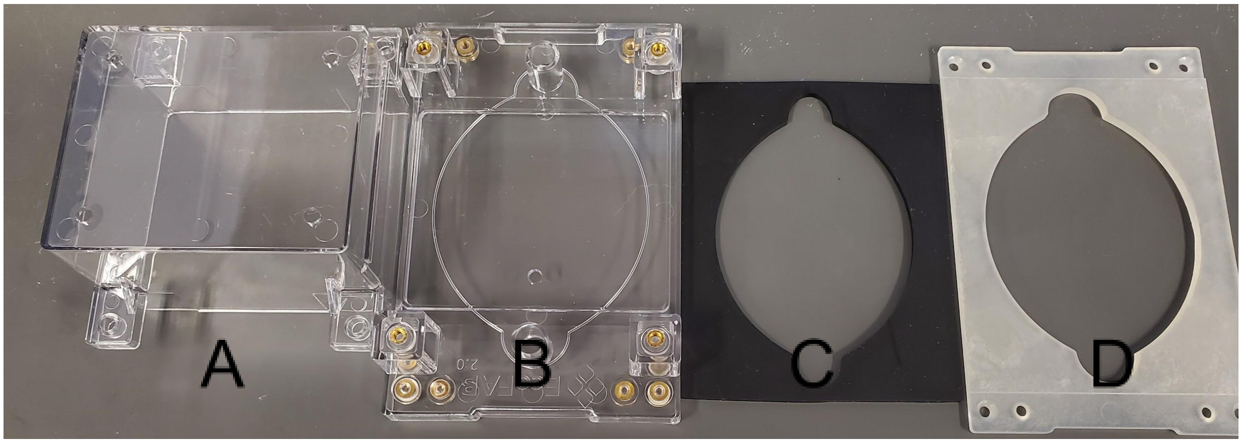


A. EcoFAB Chamber, Clear Polycarbonate (1)
B. EcoFAB Base, Polycarbonate (1)
C. Gasket, Silicone Rubber (1)
D. Backing Plate, Polycarbonate with 20% Glass Fiber (1)

2. *Additional Parts not supplied by JGI that may need to be purchased separately (quantity per device / supplier/ supplier ID/ quantity per package)*

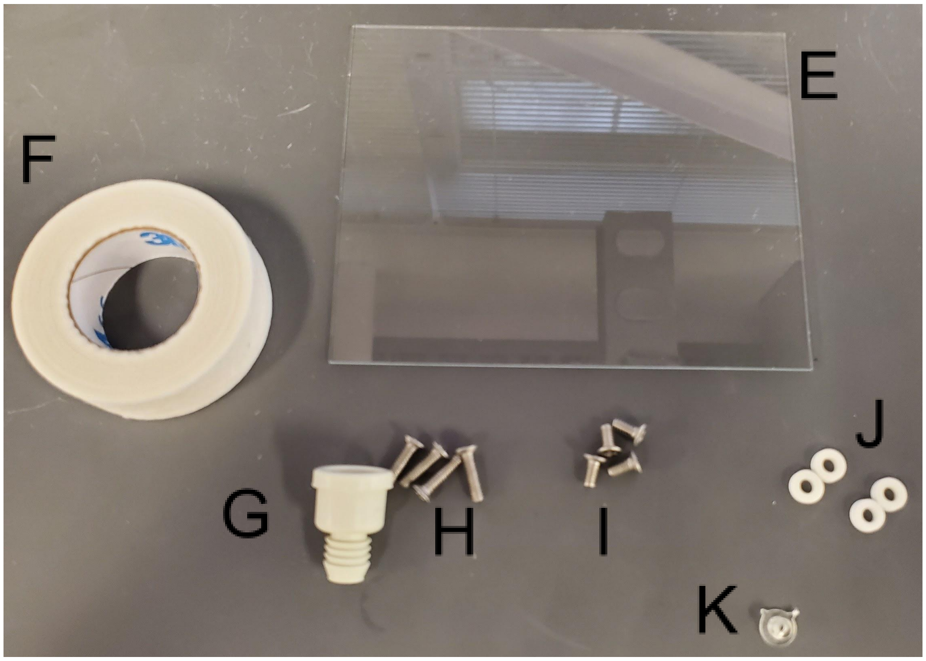


E. 4” x 3” Glass slide (1 / Ted Pella / 260230-50/ 50)
F. 3M Micropore Tape 1530-0 (a few pieces / FischerSci / 19-061655/ 24 rolls)
G. Suba-Seal 8 mm septa (1 / Sigma Aldrich / #Z167231 / 100)
H. 4/40 x 7/16” screws (4 / McMaster Carr / 91099A459 / 100)
I. 4/40 x 1/4” screws (4 / McMaster Carr / 91099A155 / 100)
J. ¼” OD EPDM Rubber Washers ( 4 / McMaster Carr / 99186A111 / 100)
K. 3D Printed Seed Inserts (1 / LBNL - optional based on request )

3. *Consumables that may be required*
    1. Serological pipette
    2. Syringes
    3. 21 G needles
    4. 0.22 micron syringe filters (PES)
    5. Sterilization pouches or aluminum foil to cover during sterilization.

**Notes on Materials**

1. The gaskets sometimes have an odor due to the manufacturing process. It is recommended to remove any plastic coatings from the gasket and then wash it with alcohol (ethanol or isopropanol) and then purified water and then allow them to sit out in a well ventilated area (e.g., biological safety cabinet or fume hood) overnight before use.
2. The EcoFAB chamber, base and backing plates can be reused however after repeated autoclaving they may warp somewhat. We have not done extensive testing to determine how many times these parts can be reused. To clean, a laboratory style dishwasher or soaps should be suitable. Alcohol may streak the polycarbonate parts.
3. The gasket will often fuse with glass slide after experiments and both parts will need to be replaced
4. One type of suba seal has been listed. There are other options available through Sigma Aldrich
5. Seed inserts are not required. However, several have been designed for smaller seeds that are 3D printed out of biologically compatible resin that can be autoclaved.
6. Alternatively to glass slides, the design is also compatible with thinner materials like 0.55 mm thick gorilla glass. These can be obtained from specialty manufacturers.
7. Instead of covering the holes on top of the chamber with micropore tape. Breathe-Easy membrane material can also be used (FisherSci).
8. The EcoFAB Base will pick up fingerprints that are visible under the microscope so wiping down the surface with a paper towel and wearing gloves is recommended.

### Sterilization

1. Most of the parts for the EcoFAB 2.0 can be sterilized by autoclaving. For most uses it is recommended to partially assemble the EcoFABs, autoclave and then finish assembly, add media and seedlings within a sterile environment (e.g., biological safety cabinet).
2. The one part that can’t be autoclaved is the micropore tape. Misting with 70% ethanol or UV may be used, but typically isn’t necessary if handled carefully.

### Assembly

1. Place the EcoFAB base upside down on a clean surface and lay the gasket down between the threaded inserts. If the gasket has matte and glossy surfaces, place the glossy surface face up so that it will be in contact with the slide in the next step.

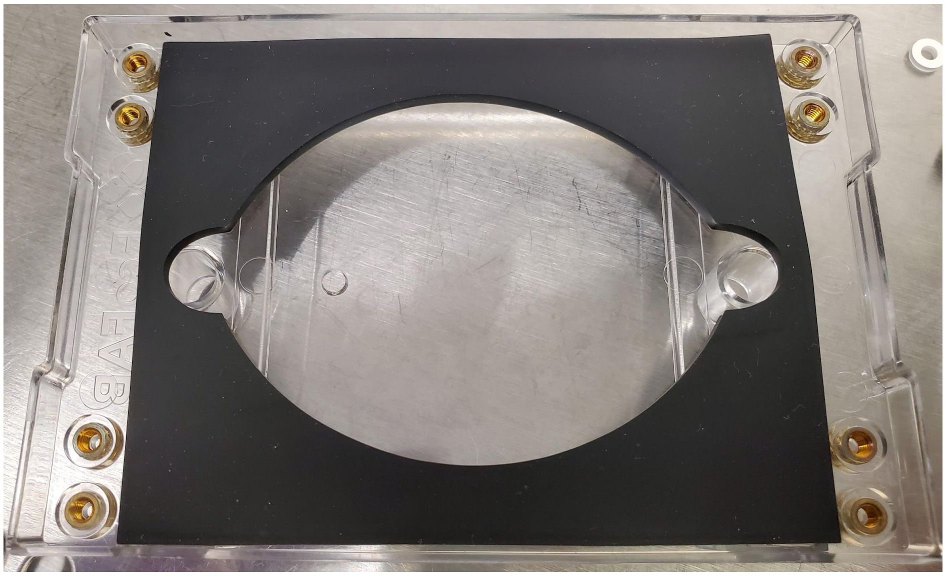

2. Place the glass slide down on top of the gasket. Make sure that the slide doesn’t overlap with the threaded inserts and covers the cutout in the gasket.

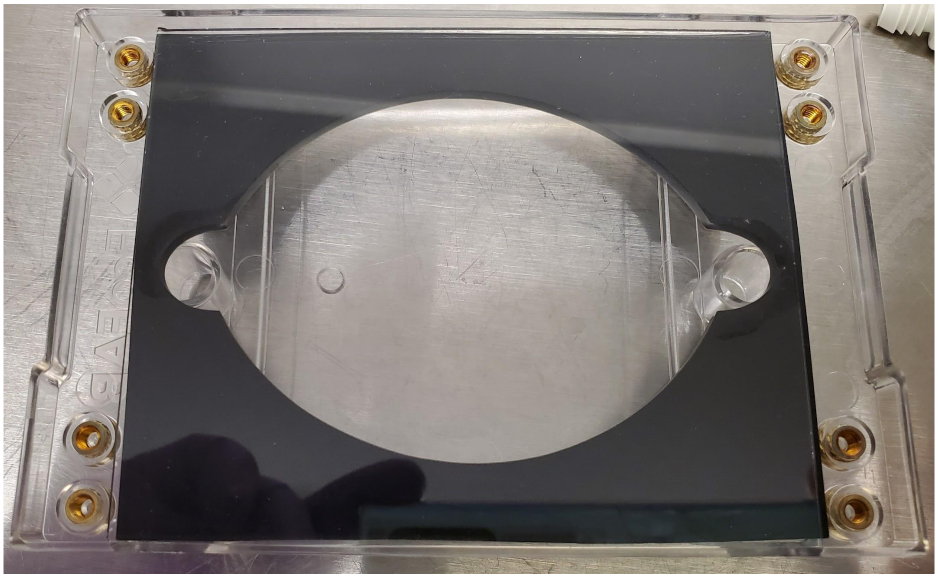

3. Place 4 washers on the four outer threaded inserts. Without the washers screwing in the backing plate will need to be done very carefully or slide will crack or the device will leak. Afterwards place the backing plate on top of everything with the protruding side facing up. (Note: the backing plate is not reversible. Usage in the other direction is for instances where a thinner slide is used. In that case the 4 internal threaded inserts will be used. You can tell this based on the indents in the backing plate screw taps.

4. Take the 4 shorter screws (¼" long) and lightly thread them into the four outer ports. If these will be sterilized do not tighten the screws enough to distort the washers or the slide is likely to crack during autoclaving. The screws should be just tight enough so that backing plate doesn't fall off when it is turned over.

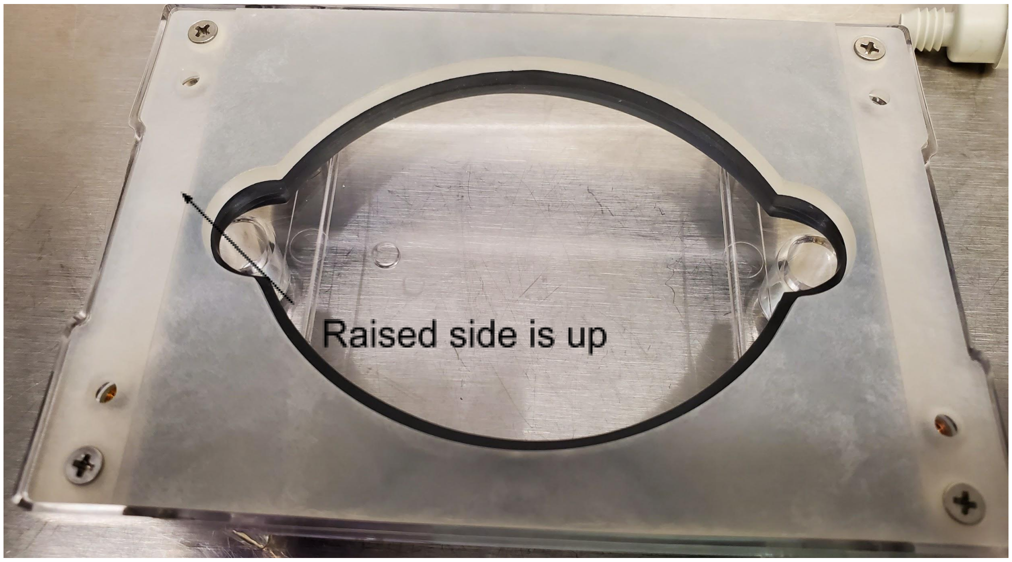


5. Place the septa and seed insert (if being used) into the EcoFAB and place the chamber on top.

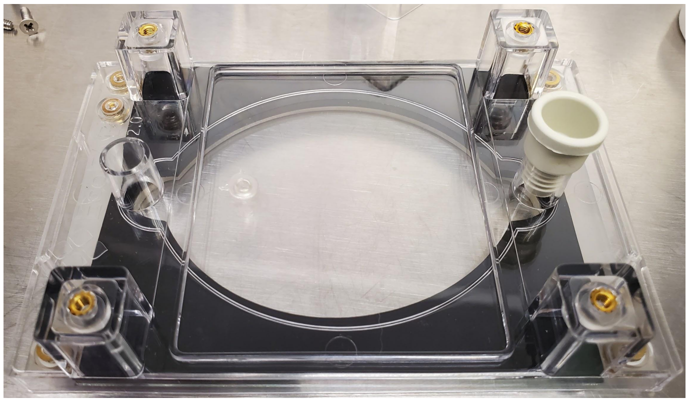

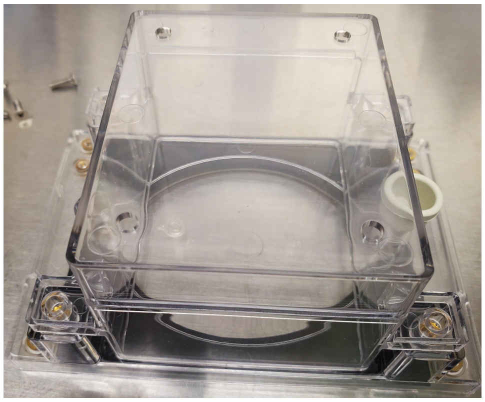

6. Place the partially assembled EcoFAB in an autoclave bag or wrap tightly in foil and autoclave to sterilize using standard conditions.

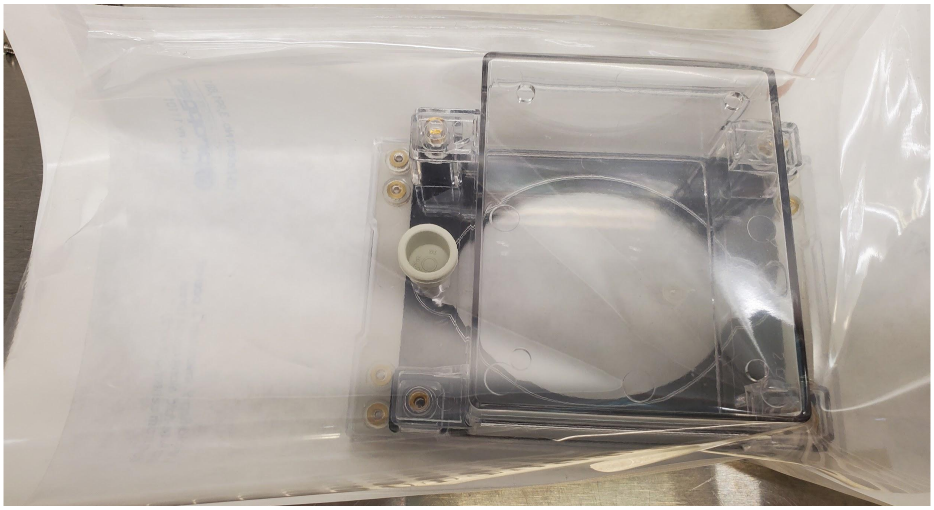

7. After sterilization, unpack the EcoFABs inside a biological safety cabinet or another sterile environment. Remove the chamber and turn the EcoFAB upside down to tighten the bottom screws. If any screws fell out during sterilization make sure the washers are aligned with the screws and the slide and gasket with the ports.

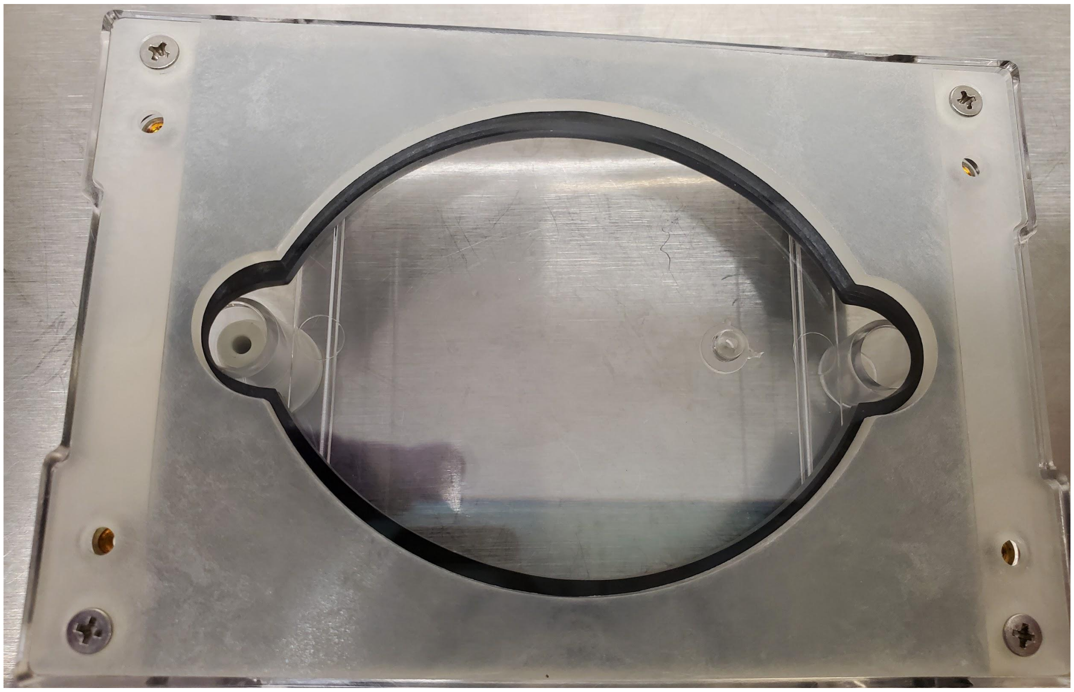

8. Place micropore tape pieces over the 4 chamber ports (or other breathable covering like Breathe-Easy membranes). Place two pieces of micropore tape over the rear sampling port to allow air exchange during filling.

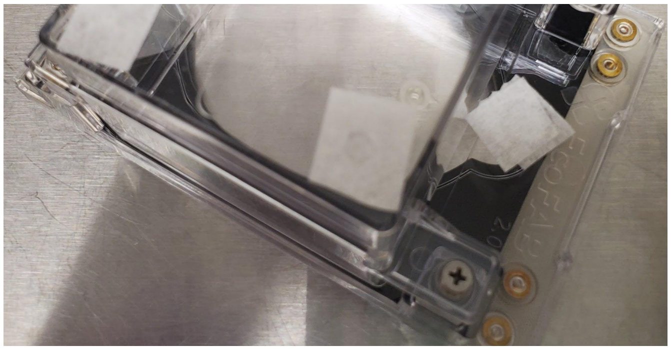

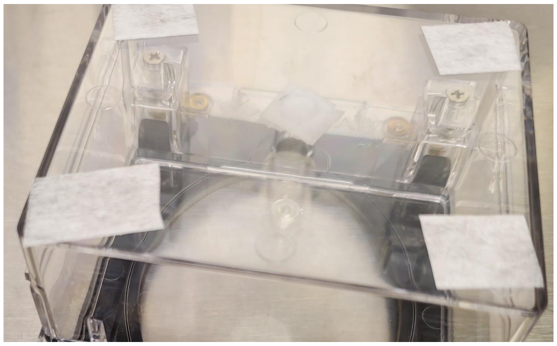

9. To fill the EcoFABs, prop the EcoFABs on an angle and fill from the bottom port either with a serological pipet with the septa removed or through the septa with a syringe and needle. Approximately 10 ml of liquid will fill the root chamber. Using sterile tweezers, place a seedling in the seed port/holder.

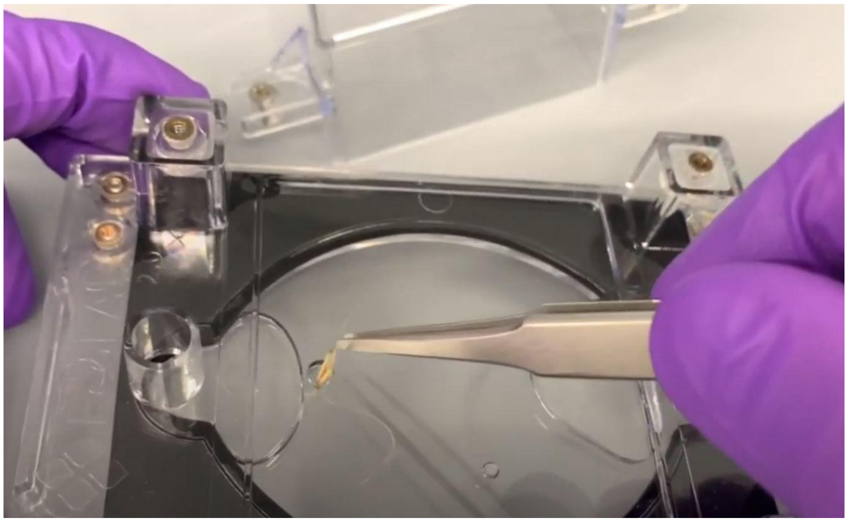

10. Screw down the top chamber and place in growth chamber.

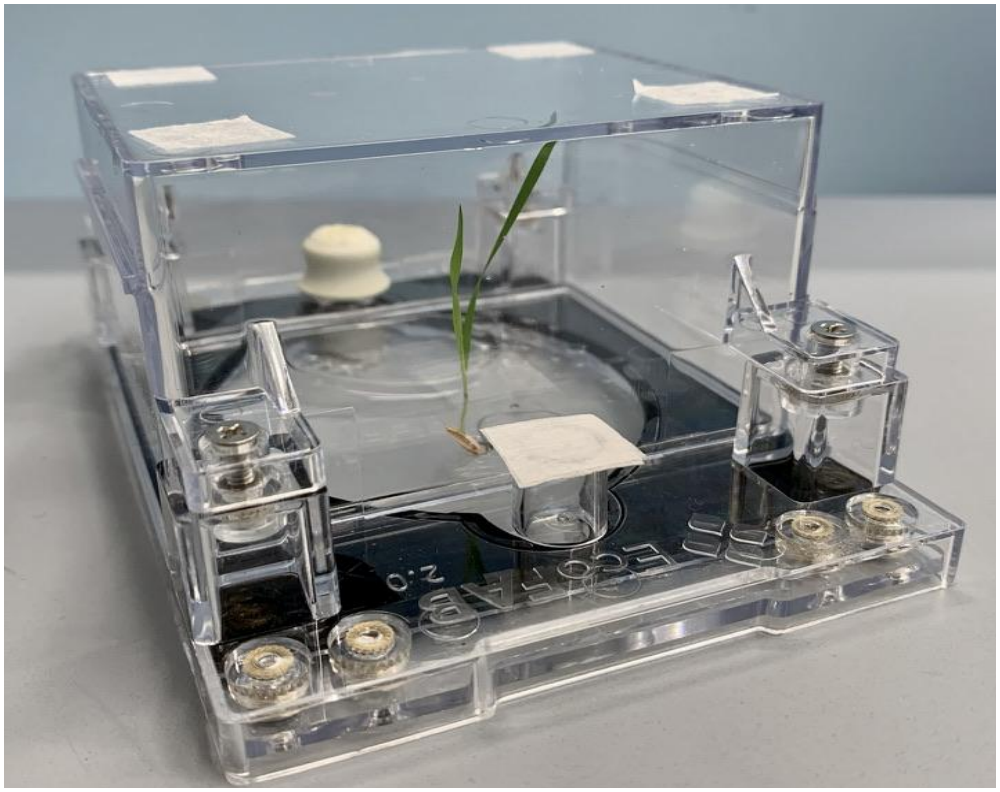
