## Supplementary material for "Reproducible growth of *Brachypodium distachyon* in fabricated ecosystems (EcoFAB 2.0) reveals that nitrogen form and starvation modulate root exudation": Figures S1-S5

**This PDF file includes:**

Figs. S1 to S5

**Other Supplementary Materials for this manuscript include the following:**

Protocol S1
Tables S1 to S6
Files S1 to S2

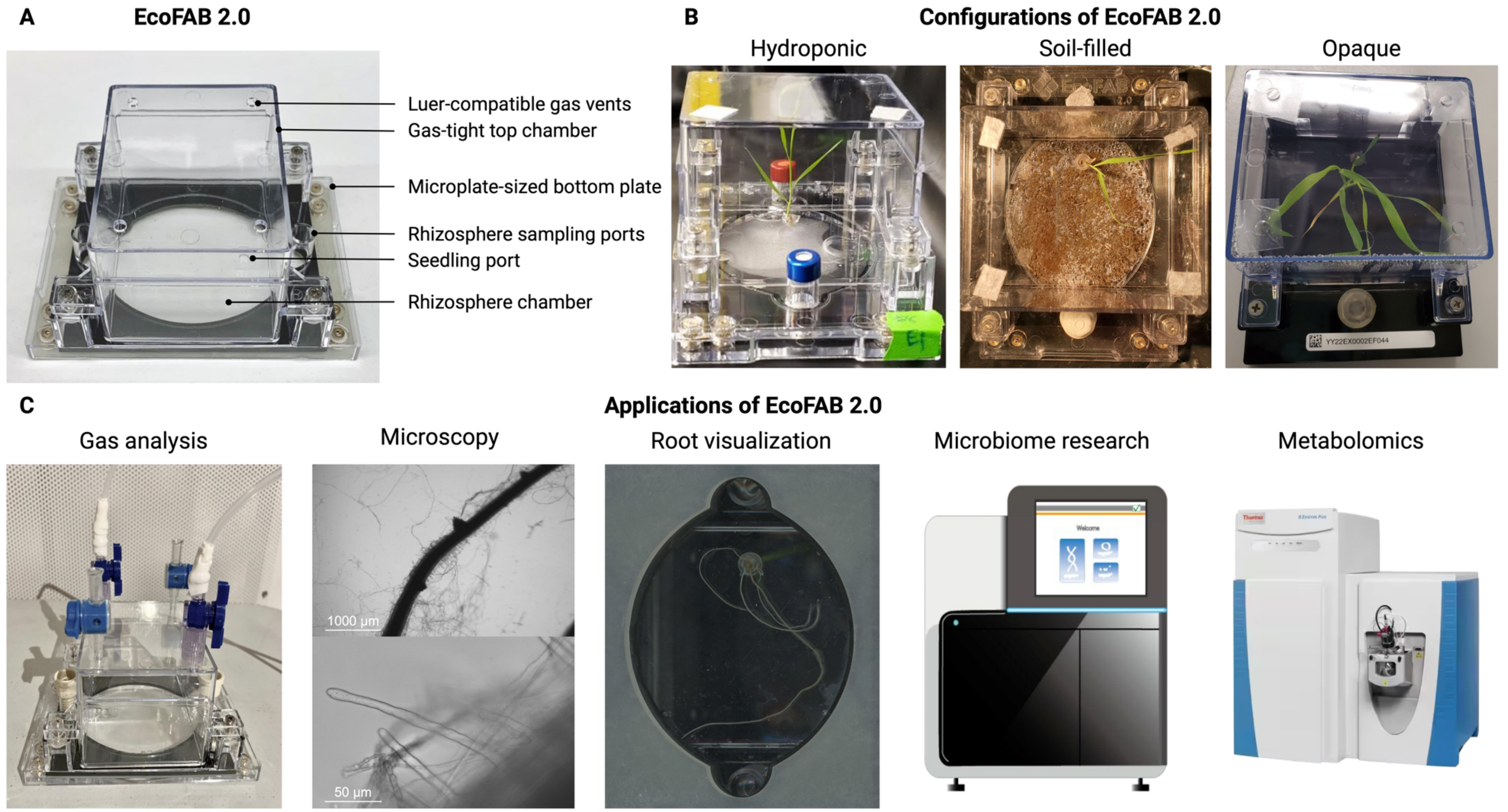


**Fig. S1: EcoFAB 2.0 devices can accommodate a wide diversity of plant research.** (a) Empty EcoFAB 2.0 and its main functional parts. (b) Configurations of EcoFAB 2.0 for *B. distachyon* growth include (from left): EcoFAB 2.0 filled with liquid medium for hydroponic plant growth (used in this study), soil-filled setup, and a version of EcoFAB 2.0 with opaque black root chamber that allows light-sensitive rhizosphere experiments. (c) Examples of possible applications include (from left): gas analysis, microscopy of roots (2X and 40X magnification), root system visualization, study of microbiomes by 16S sequencing, and metabolomics on growth medium (used in this study).

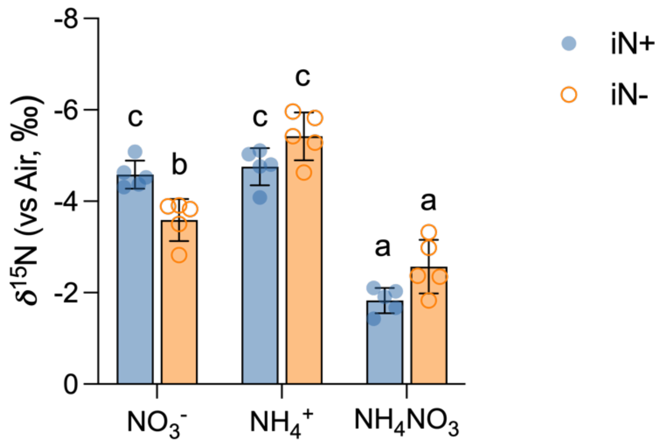


**Fig. S2: Shoots N stable isotope ratios.** Bars show mean ±SD of iN+ of control (blue) and iN- (orange) plants. Different letters indicate statistically significant differences (two-way ANOVA with post hoc Tukey's HSD test; $n = 5$; $p \leq 0.05$).

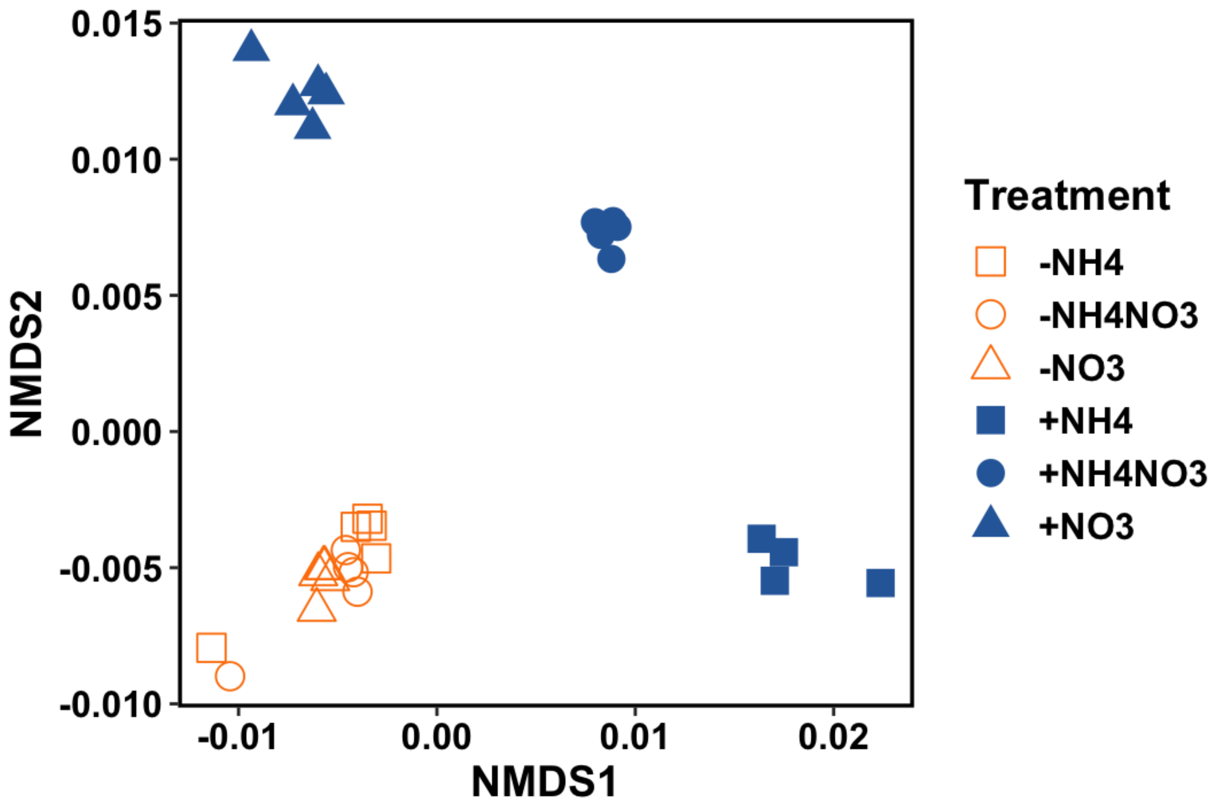


**Fig. S3: NMDS plot for *B. distachyon* root exudate features using raw peak height data (filtered features, *n* = 2065).** Each biological replicate across iN treatments is represented with a symbol. The Blue symbols represent iN+ treatments, while the orange symbols represent plants grown in iN- medium.

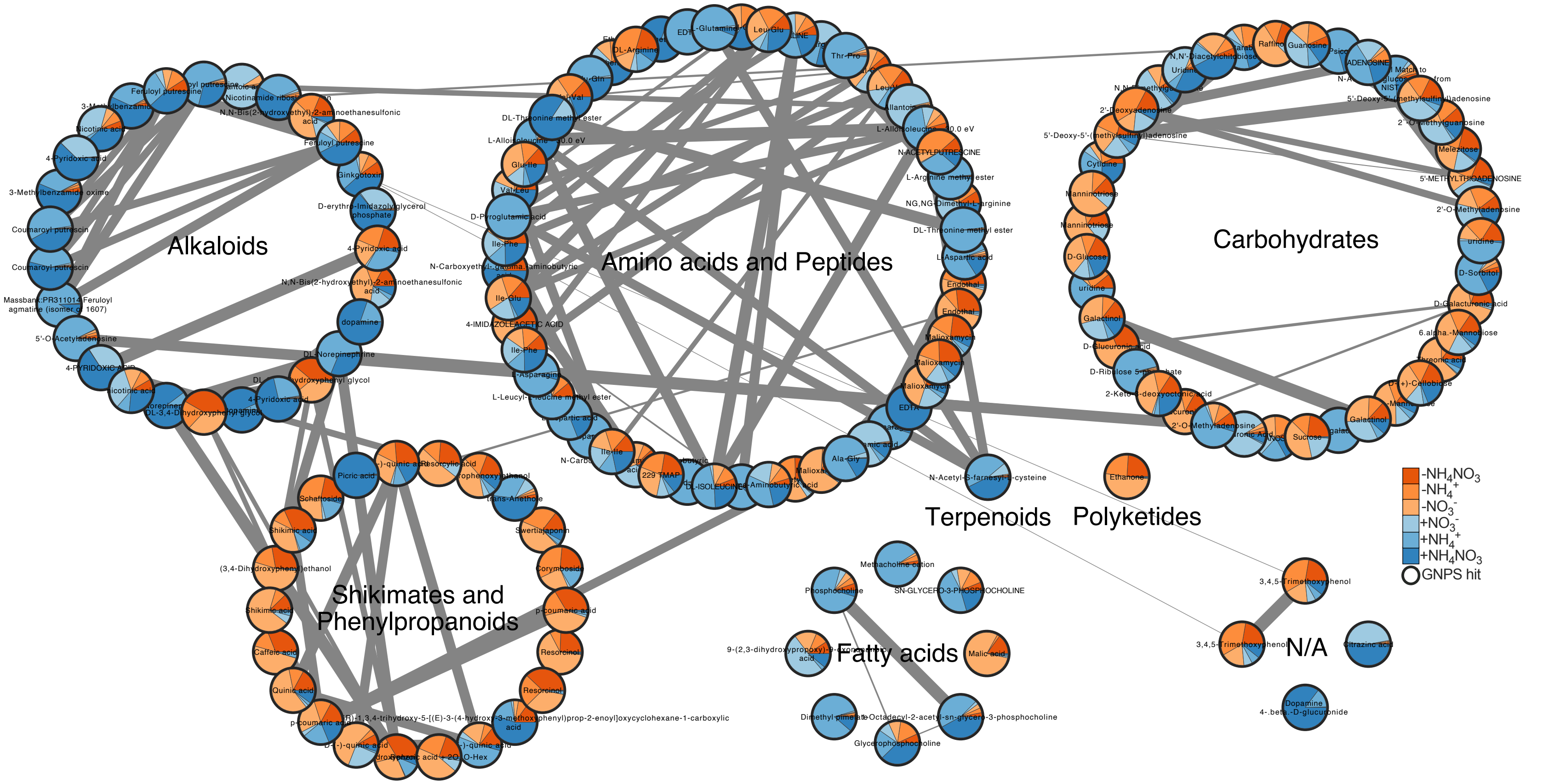


**Fig. S4: Sub-network of grouped annotated features.** The features were grouped into biosynthetic pathways by NPClassifier. Pie charts show relative mean peak heights for individual iN- treatments (shades of orange) and iN+ treatments (shades of blue). The black borders indicate GNPS annotation with MQScore > 0.7 (*n*=155). The network shows merged features from positive and negative polarities of polar metabolite analysis in root exudates of *B. distachyon* Bd21-3 at week 5.

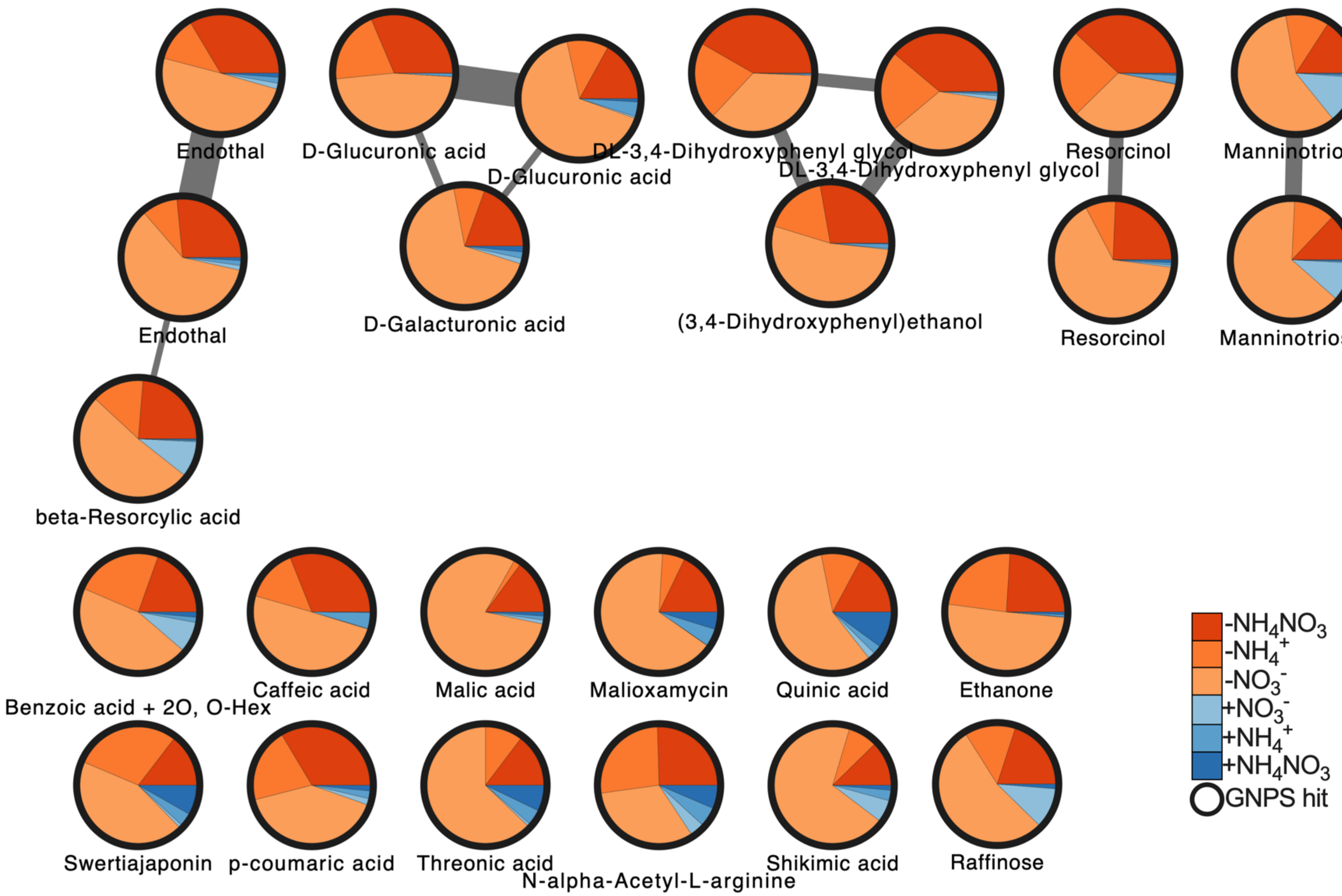


**Fig. S5. Sub-network of annotated features abundant in N-deficient root exudates.** The network shows features significantly increasing > 5-fold in iN- treatments ($n$=15) relative to the iN+ treatments ($n$=14) (t-test, $p$-value ≤ 0.5). Pie charts show relative mean peak heights for individual iN- treatments (shades of orange) and iN+ treatments (shades of blue). The black borders indicate GNPS annotation with MQScore > 0.7. The network shows merged features from positive and negative polarities of polar metabolite analysis in root exudates of *B. distachyon* Bd21-3 at week 5.
